# Mechanistic insights into zearalenone-accelerated colorectal cancer in mice using integrative multi-omics approaches

**DOI:** 10.1101/2022.09.21.508814

**Authors:** Emily Kwun Kwan Lo, Xiuwan Wang, Pui-Kei Lee, Ho-Ching Wong, Jetty Chung-Yung Lee, Carlos Gómez-Gallego, Danyue Zhao, Hani El-Nezami, Jun Li

## Abstract

Zearalenone (ZEA), a secondary metabolite of *Fusarium* fungi found in cereal-based foods, promotes the growth of colon, breast, and prostate cancer cells *in vitro*. However, the lack of animal studies hinders a deeper mechanistic understanding of the cancer-promoting effects of ZEA. This study aimed to determine the effect of ZEA on colon cancer progression and its underlying mechanisms. Through integrative analyses of transcriptomics, metabolomics, metagenomics, and host phenotypes, we investigated the impact of a 4-week ZEA intervention on colorectal cancer in xenograft mice. Our results showed a twofold increase in tumor weight with the 4-week ZEA intervention. ZEA exposure significantly increased the mRNA and protein levels of BEST4, DGKB, and Ki67 and the phosphorylation levels of ERK1/2 and AKT. Serum metabolomic analysis revealed that the levels of amino acids, including histidine, arginine, citrulline, and glycine, decreased significantly in the ZEA group. Furthermore, ZEA lowered the alpha diversity of the gut microbiota and reduced the abundance of nine genera, including *Tuzzerella* and *Rikenella*. Further association analysis indicated that *Tuzzerella* was negatively associated with the expression of BEST4 and DGKB genes, serum uric acid levels, and tumor weight. Additionally, circulatory hippuric acid levels positively correlated with tumor weight and the expression of oncogenic genes, including ROBO3, JAK3, and BEST4. Altogether, our results indicated that ZEA promotes colon cancer progression by enhancing the BEST4/AKT/ERK1/2 pathway, lowering circulatory amino acid concentrations, altering gut microbiota composition, and suppressing short chain fatty acids production.

## Introduction

Colorectal cancer (CRC) is one of the most common cancers and the third leading cause of cancer-related death worldwide [1]; however, the pathogenesis of CRC remains to be elucidated. Recently, increasing evidence has suggested that dysbiosis of the gut microbiome may contribute to CRC development [2–4]. Exposure to exogenous estrogen is an environmental hazard contributing to CRC progression by binding to estrogen receptors [5].

Zearalenone (ZEA) is a mycotoxin produced by *Fusarium* spp. and is commonly found in the diet, mainly in cereals and grains. Owing to its structural similarity to estrogen, ZEA promotes *in vitro* cancer cell growth and progression in hormone-dependent cancers, including breast [6] and prostate cancers [7]. A recent epidemiological study has reported a link between increased levels of urinary ZEA metabolites and breast cancer development [8]. Moreover, studies using colon cancer cell lines have shown that ZEA accelerates colon cancer cell growth and metastasis [9, 10]. However, these studies disregarded the three-dimensional growth, tumors microenvironment, and immune regulatory role of the gut microbiota in living animals [11]. Because ZEA is mainly digested and absorbed through the gastrointestinal tract, the gut microbiota is directly exposed to ZEA, which potentially regulates its metabolism and contributes to carcinogenesis. Various studies have explored the impact of ZEA on the composition of the gut microbiota and indicated that ZEA might disrupts the composition and function of the gut microbiota [12–14]. However, these studies were conducted exclusively on healthy individuals.

Given that ZEA is commonly found in the human diet and that its impact can be persistent [15], an in-depth mechanistic understanding of the potential contribution of ZEA exposure to colon cancer development is warranted. In this study, we comprehensively evaluated the effects of ZEA exposure on the development and progression of CRC in a colon cancer xenograft mouse model. This was achieved through an integrated analysis of the tumor transcriptome, targeted and untargeted metabolomics, and 16S rDNA sequencing. Furthermore, we investigated the mechanisms underlying the accelerated tumor growth and identified the key transcriptional signatures, gut microbes, and metabolites that potentially contribute to ZEA-induced colon cancer development.

## Materials and Methods

### Animal study

Nude BALB/cAnN mice were purchased from the Laboratory Animal Unit at the University of Hong Kong. Five-week-old male mice were housed in individually ventilated cages in a sterile environment with regulated temperature (23–24°C) and relative humidity (60–70%) on a 12-hour light cycle. Mice were fed a phytoestrogen-free diet (Open Standard diet, D11112201; Research Diets Inc., New Brunswick, NJ, USA) with water provided *ad libitum*. After one week of acclimatization, the mice were injected in the right flank with 1x10^6^ SW480 human colon cancer cells mixed with PBS or Matrigel (1:1). When the tumor size reached 70–100 mm^3^; the mice were randomly divided into two treatment groups (three independent experiments with n = 3 per group; n = 9 per group in total): control [oral gavage vehicle (olive oil) thrice weekly] and ZEA [0.5 mg/kg body weight [BW] in olive oil thrice weekly) (Fig. 1A). The dosage in this study was chosen based on an animal study in which 1 mg/kg BW ZEA promoted DNA adduct formation in healthy mice, and the reported levels of ZEA in grain-based food in humans (3.0–33.0 µg/kg bw/day) [16, 17], in which 33.0 µg/kg bw/day was converted through body surface area conversion from humans to mice [18]. Tumor size was measured as previously described, with some modifications [19]. Tumor sizes were measured using an electric caliper three times weekly. Tumor volumes were calculated based on the formula: (volume [mm^3^]LJ= π x length× width^2^/6). Length represents the longest tumor diameter (mm), and width represents the perpendicular diameter (mm). BW and food consumption were recorded three times per week throughout the study. After 28 days, the mice were sacrificed using pentobarbital (250 mg/kg, intraperitoneal injection), and the tumor and cecal contents were removed, weighed, and snap-frozen in liquid nitrogen. Animal handling was approved by the Committee on the Use of Live Animals in Teaching and Research (CULATR No. 4785-18) at the University of Hong Kong.

**Fig. 1.**
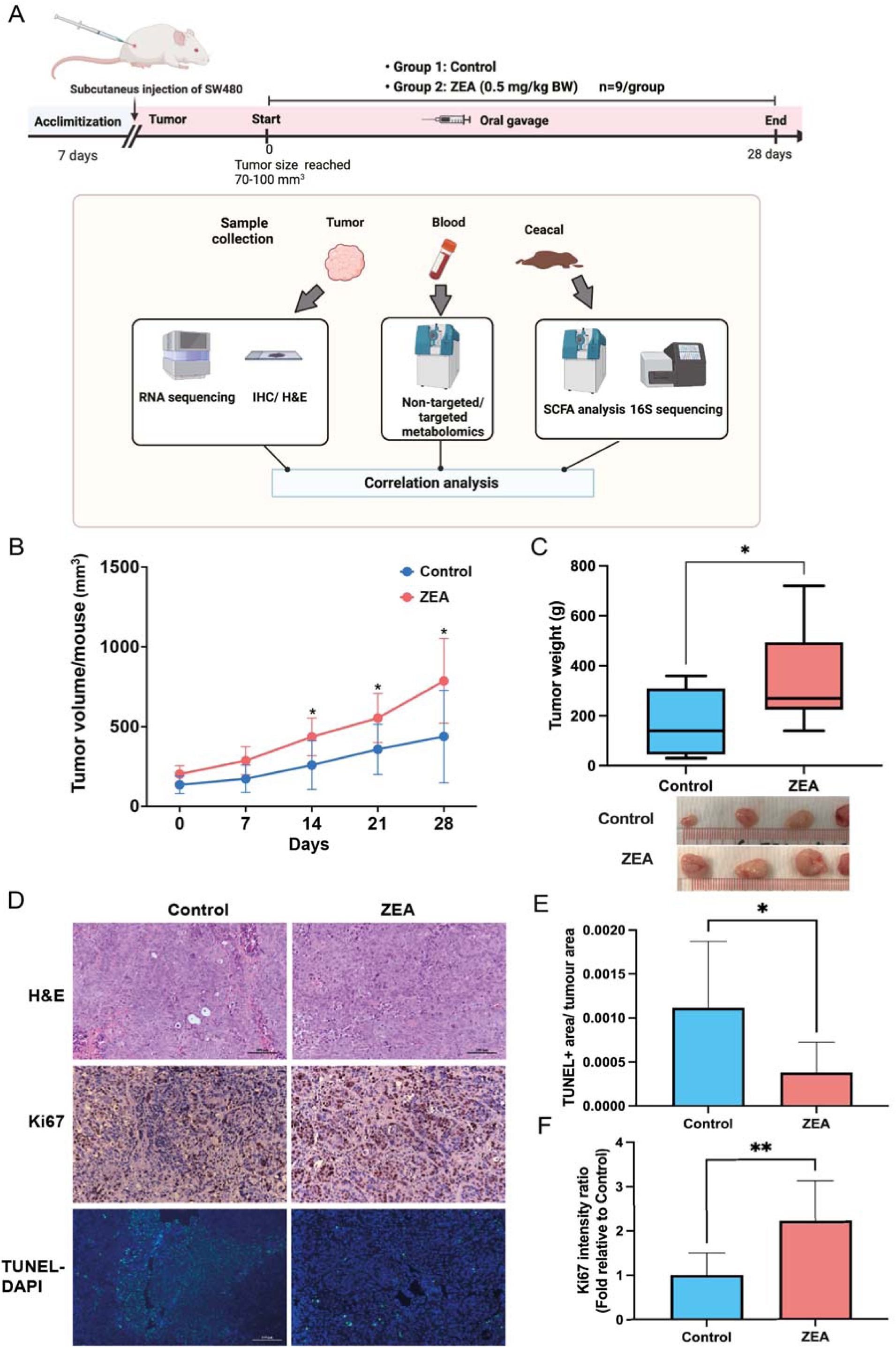
Zearalenone (ZEA) accelerated tumor growth in a colon cancer xenograft mouse model. (A) Study design. Male 6-week-old Balb/c nude mice (n = 9) were treated with vehicle (olive oil) or ZEA (0.5 mg/kg BW) thrice per week after subcutaneous injection of the human colon cancer cell line SW480. (B) Tumor volume variation during 28 days of monitoring, n = 9. (C) Weight of tumors on day 28 and representative photograph of excised tumors imaged after ZEA treatment. (D) Representative microscopic photographs of Hematoxylin-eosin (H & E) staining, Ki67 immunohistochemistry (IHC) staining, terminal deoxynucleotidyl transferase dUTP nick end labeling (TUNEL) staining (200x). (E) Quantification of TUNEL labelling was quantified as the total pixel area of the tumor. (F) while the expression of Ki67 was quantified as the relative Ki67-positive nuclei compared to the control. Signal normalized to the total pixels of the tumor (n=6-7). * *p* < 0.05, \*\**p* < 0.01, compared to control.

### RNA extraction and RNA-seq

According to the manufacturer’s instructions, RNA was isolated from the tumor tissue using an RNAspin Mini kit (GE Healthcare, UK). After passing the quality check, the samples (six randomly selected individuals per group) were fragmented and cDNA was synthesized through reverse transcription. The final cDNA library was then built using Novogene (Novogene, China) with the following procedures: purification, terminal repair, A-tailing, ligation of sequencing adapters, size selection, and PCR enrichment. Finally, reads were generated using an Illumina HiSeq 2500.

### Quality control of RNA-Seq reads and quantification of gene expression

All raw reads were mapped to the GRCm38 (Genome Reference Consortium Mouse Build 38, NCBI BioProject Accession: PRJNA20689) genome using BWA v0.7.17, and reads with > 95% mapping coverage were removed. Ribosomal RNA (rRNA) reads were filtered through SortMeRNA [20]. Raw reads were further processed for quality control using the following procedures [21] (available at https://github.com/TingtZHENG/metagenomics/blob/master/scripts/fqc.pl): (i) Illumina primers/adaptors/linker sequences were removed; (ii) paired-end reads with 25 bp consecutively exact matches from both ends were removed to avoid PCR duplicates, and (iii) terminal regions with continuous Phred-based quality of less than 20 were removed. The remaining high-quality reads were aligned using TopHat2 v2.1.1, employing the GRCh38 (Genome Reference Consortium Human Build 38, NCBI BioProject Accession: PRJNA31257) assembly as the human reference genome. After that, alignment outputs from TopHat2 were formatted and sorted using SAMtools v1.9. SAMtools outputs were used to perform gene counts using the htseq-count function of the HTseq framework v0.11.2. Additionally, RSEM (v1.3.1) was used to estimate fragments per kilobase of transcript per million mapped reads (FPKM) based on the alignment results. Based on the read count of the genes for each sample, we first applied a centered log-ratio transformation.

### Analysis of differentially expressed genes and enriched pathways

Subsequently, differentially expressed genes were identified using differential gene expression analysis based on the negative binomial distribution (DESeq2) v1.34.0, which was used to calculate the significance of the gene expression profile alterations in the comparison groups. The adjusted *p*-value < 0.1 were used as the cutoff value for significantly differentially expressed genes. Next, differentially expressed pathways were identified by Generally Applicable Gene-set Enrichment (GAGE, v2.44.0) analysis using gene sets derived from Kyoto Encyclopedia of Genes and Genomes (KEGG) pathways. An adjusted *p*-value < 0.1 was used to define significantly differentially expressed pathways in the comparison groups. Additionally, gene set enrichment analysis (GSEA, v4.0.3) was performed on gene expression profiles using gene sets from the Molecular Signatures Database (MSigDB, v7.1): (1) Cancer Hallmarks, (2) Oncogenic Signatures, and (3) Chemical and Genetic Perturbations. |NES score| > 1 and FDR < 0.25 were used as the cutoff values for significantly enriched gene sets.

### Hematoxylin-eosin (H & E) staining, immunohistochemistry (IHC), and TUNEL assay

Part of the tumor tissue was washed, fixed immediately with formalin, and embedded in paraffin for sectioning into 5 mm slices. The sections were then deparaffinized and subjected to H & E, IHC, or terminal deoxynucleotidyl transferase dUTP nick end labeling (TUNEL) fluorescent staining. According to the manufacturer’s recommendations, H & E staining was performed using an H & E kit (Baso, Zhuhai, China). The TUNEL assay was performed using the In Situ Cell Death Detection Kit (Roche Diagnostics) according to the manufacturer’s instructions, with some modifications [22]. In brief, the slides were incubated in 0.1 M citrate buffer in a 70°C water bath for 30 min and then blocked with CAS block (Thermo Fisher, USA). The slides were washed and then incubated with a TUNEL reaction mixture for 60 min at 37°C. For IHC, sections were rehydrated in graded alcohol and distilled water. It was then boiled in Tris-EDTA, followed by quenching of endogenous peroxidase activity using 3% H_2_O_2_. The sections were immersed in CAS-Bloc Histochemical Reagent (Thermo Fisher, USA) and incubated with primary antibodies against pERK1/2, pAKT, Ki67, BEST4, (1:100, Abcam) and DGKB (Invitrogen, USA) at 4°C overnight. They were then washed thoroughly and incubated with anti-mouse, anti-rabbit horseradish peroxidase-conjugated secondary antibodies (Bio-Rad) or anti-goat horseradish peroxidase-conjugated secondary antibodies (Bio-Rad). Positive signals were visualized using 3,3’-diaminobenzidine (Abcam). Nuclei were counterstained with hematoxylin. Expression in IHC images was quantified by NIH ImageJ v1.50, as previously described [23].

### Non-targeted metabolomics analysis

One hundred microliters of serum sample were added to 1 ml ice-cold 100% methanol, spiked with 4-chloro-phenylalanine as the internal standard, at a final concentration of 100 ng/ml. The mixture was vortexed for 30 s and kept at -20°C for 20 min. The cold mixture was centrifuged at 17,000 x g at 4°C for 15 min. The supernatant was collected and dried under gentle nitrogen flow. The dried samples were reconstituted with 100 µL 60% methanol and sonicated for 1 min before centrifugation at 17,000 x g for 10 min. The supernatant collected was subjected to LC-MS/MS analysis using Agilent 6540 UHPLC-QTOF-MS/MS (Agilent Technologies, Santa Clara, CA, USA) with a Waters ACQUITY HSS T3 column, 2.1 mm x 100 mm, 1.7 μm (Milford, MA). The column temperature was set at 40°C, and the injection volumes for positive and negative modes were set at 10 µL and 15 µL, respectively. The mobile phase consisted of water containing 0.1% formic acid (A) and acetonitrile containing 0.1% formic acid (B) at a flow rate of 0.3 ml/min. The gradient (B%) started at 1% from 0 to 1 min, increased to 99% from 1 to 10 min, held at 99% until 13 min, returned to 1% at 13.5 min and maintained until 16 min. The MS parameters were as follows: capillary voltage 4000 V, nozzle voltage 1500 V, skimmer voltage 65 V, drying gas temperature 300°C, sheath gas temperature 320°C, fragmentor voltage 140 V, drying gas flow rate 8 L/min, sheath gas flow rate 11 L/min, and nebulizer pressure 40 psi. Metabolites were identified by comparing the data to the HMDB and METLIN databases using Progenesis QI software version 3.0 (Nonlinear Dynamics, Newcastle, UK).

### Targeted metabolomics for amino acid profiling

Serum samples (50 μL) (n = 6 per group) were mixed with 50 μL of an internal standard (Kairos Amino Acid Internal Standard Set, Waters, USA) and water according to the manufacturer’s instructions. The mixture was centrifuged at 9000 x g for 15 min. The mixture (10 μL) was then mixed with 70 μL borate buffer and 10 μL of AccQ•Tag reagent. After vortexing for 5 s, the mixture was allowed to stand at room temperature for 1 min and heat for 10 min at 55°C. The mixture was then analyzed using Agilent 6460 UHPLC-QqQ-MS/MS (Agilent Technologies, Santa Clara, CA, USA) with a Waters Cortecs C18 column, 2.1 mm x 150 mm,

1.6 μm (Milford, MA). For chromatographic separation, a binary mobile phase system was used, including 0.1% formic acid in water (A) and 0.1% formic acid in acetonitrile (B), at a flow rate of 0.5 ml/min. The gradient elution program (B%) for each run started at 1%, lasted for 1 min, and then increased to 13% at 2 min, 13% to 15% from 2 to 5.5 min, 15% to 95% from 5.5 to 6.5 min, kept at 95% until 7.5 min, then decreased to 1% from 7.5 to 7.6 min and maintained at 1% until 9 min. The column temperature was 55°C, and the injection volume was 2 μL. Data processing was performed using Agilent MassHunter Workstation software (version B.06.00).

### Metabolomics data analysis

Pathway analysis was performed to integrate the targeted amino acid results using the Ingenuity Pathway Analysis Software (IPA) version 70750971 (QIAGEN, Redwood City, USA). Metabolites strongly correlated with tumor weight were identified using Spearman’s rank correlation analysis. The |correlation| > 0.6 and *p-value* < 0.05 were used as the cutoff values of statistical significance.

### Total microbial DNA extraction and 16S rDNA gene quantitative analysis

The DNA content of the cecal samples was extracted using a DNeasy PowerSoil Kit (Qiagen, Germany) according to the manufacturer’s instructions. The quality of the extracted DNA samples was examined by 1% agarose gel electrophoresis. The DNA samples (n = 6 per group) were sent to Novogene Co., Ltd. (Beijing, China) for sequencing. After a quality check, the hypervariable region V3-V4 (341F and 534R) of the 16S rDNA was amplified. PCR was performed using Phusion High-Fidelity PCR Master Mix (New English Biolabs, England). PCR products were quantified using 2% agarose gel electrophoresis. Products between 400 and 450 bp were extracted using a Qiagen Gel Extraction Kit (Qiagen, Germany). The samples were quantified using a Qubit and prepared for sequence library generation using the NEBNext Ultra DNA Library Prep Kit. The resulting library was sequenced using an Illumina HiSeq 2500 platform. The R package dada2 v1.22.0 was used to perform read quality control and resolve amplicon sequence variants (ASVs). Thirty low-quality 3’ bases of the forward and reverse reads were removed after inspection of the QC plots. Reads with any Ns or maximum expected error > 2 after truncation and mapping to phiX were also removed. Error estimation was performed for all samples (pooled) after de-replication. Subsequently, dada2 merged the overlapping forward and reverse reads, and the chimeric sequences were removed. Following chimaera removal, taxonomy was assigned independently to the SILVA v138.1 reference database. Shannon diversity at the genus level was calculated using the R package vegan v2.6.2. Alpha diversity indices were compared between the experimental and control groups using an independent-sample t-test. The R packages vegan and compositions v2.0.4 [24] were used to calculate beta diversity based on the robust Aitchison distance. The Wilcoxon rank-sum test was used to evaluate whether there was a significant difference in the pairwise distance within groups between the comparison groups. We then evaluated beta diversity using multiple dimension scale (MDS) ordinations with vegans. Permutational multivariate analysis of variance (PERMANOVA) of the distance matrices between groups was performed to calculate R^2^ (0 = no difference; 1 = maximum dissimilarity) and determine group differences. Differential abundance of taxa at the genus level between comparison groups was determined using the Analysis of Compositions of Microbiomes with Bias Correction (ANCOM-BC) implemented in the R package ANCOMBC v1.4.0 [25]. An adjusted *p-value* < 0.05 was used as the cutoff to define significantly differentially abundant genera between groups.

### Gene expression, microbiota, and metabolome integrative analysis

We performed an integrative analysis to reveal associations between multi-omics and phenotypic data. Spearman’s correlation coefficients were calculated between the differentially expressed genes, differentially abundant microbial taxa and differentially abundant metabolites between the experimental, and control groups. Statistical significance was set at FDR < 0.05 after Bonferroni correction, and a strong correlation was defined as |correlation| > 0.8. Furthermore, we built significantly strong correlations in a visualization network using Cytoscape software v3.9.1.

### Short chain fatty acids (SCFAs) analysis of cecal contents

The concentration of SCFAs in the caecum was measured by gas chromatography-mass spectrometry (GC-MS), as previously described, with modifications [26, 27]. In brief, the cecal content samples were homogenized using a blade homogenizer (T25, ULTRA-TUREAX, IKA) in an extraction buffer containing 0.005 M sodium hydroxide with internal standard (10 µg/ml acetic acid-d4). The homogenates were centrifuged at 13,200 x g for 20 min at 4°C. The supernatant was mixed with 0.5 ml of 1-propanol/pyridine (3:2, v/v) and 0.1 ml propyl chloroformate. The mixture was vortexed for 1 min and incubated at 60°C to derivatize SCFA. The derivatized samples were mixed with hexane (0.5 ml) and centrifuged at 2000 x g for 4 min. After that, 400 µl of the upper layer was transferred to a glass vial for GC–MS analysis (Agilent 6890N-5973 GCLJMS, USA) set according to Zheng et al. [26] report. The concentration of SCFAs was quantified using calibration curves constructed using acetic acid, propionic acid, and the butyric acid response ratio against acetic acid-d4.

### Statistical analysis

Data are presented as the means ± SDs. Comparisons of experimental results were performed using an independent-sample t-test on GraphPad Prism 9.0 (GraphPad Software, San Diego, CA, USA) unless specified otherwise. *P-values* < 0.05 were considered statistically significant. Heatmap visualization was performed using the R package heatmap v1.0.12, and other figures were generated using the R package ggplot2 v3.3.5.

## Data Availability Statement

Transcriptome and 16S rDNA data were archived in the National Center for Biotechnology Information (NCBI) Short Read Archive (SRA): PRJNA875586.

## Results

### Body weight and feed intake were unaffected by ZEA

ZEA administration was well tolerated by the mice, with a steady increase in body weight (Fig. S1A) and the absence of reduced motor function. There were no significant differences in the weights of the major organs, including the liver, spleen, lungs, heart, or kidneys (Fig. S1C) among the treatment groups. Interestingly, the body weight of the ZEA group peaked after 24 days of treatment and decreased thereafter; it is anticipated that the increase in tumor burden affected them. To determine whether the decrease in body weight was due to reduced feed consumption, we analyzed the food intake throughout the treatment. The results revealed no significant differences in feed consumption (Fig. S1B). In summary, these results indicated that body weight and feed intake were unaffected by ZEA intervention.

### ZEA promoted tumor growth

We observed a nearly linear increase in the tumor volume over time in both the control (vehicle) and ZEA groups (Fig. 1B). The tumor volume was significantly larger than that in the control group (*p* < 0.001) at the end of the intervention (Fig. 1B). The tumor weight in the ZEA group tripled (*p* = 0.036 with an independent sample t-test) compared to that in the control group (Fig. 1C). The TUNEL assay showed stronger inhibition of tumor apoptosis by ZEA than by the control (Fig. 1D–E). Consistently, IHC staining showed that Ki67-positive cells in tumor tissues were significantly increased twofold (*p* = 0.002 with independent sample t-test) in the ZEA group compared with the control (Fig. 1D and F). We showed that ZEA reduced cancer cell death and promoted tumor cell proliferation.

### Changes in gene expression associated with ZEA-induced tumor growth

DESeq2 identified 12 significantly upregulated and nine downregulated genes (Table 1 and Fig. 2A) in the ZEA group compared to those in the control group. Based on the overall expression profile across different samples, these differentially expressed genes (DEGs) were grouped into three clusters (Fig. S2). Cluster 3 contained most of the significantly upregulated genes, including DGKB, EBF2, PTGIR, and BEST4. An over 20-fold increase in expression (log_2_ fold change = 4.37) was observed in DGKB (Table 1), which was previously reported to be a gene signature for the prognostic prediction of CRC [28]. Another upregulated gene worth mentioning is BEST4, which was previously reported to be overexpressed in patients with CRC [29].

**Fig. 2.**
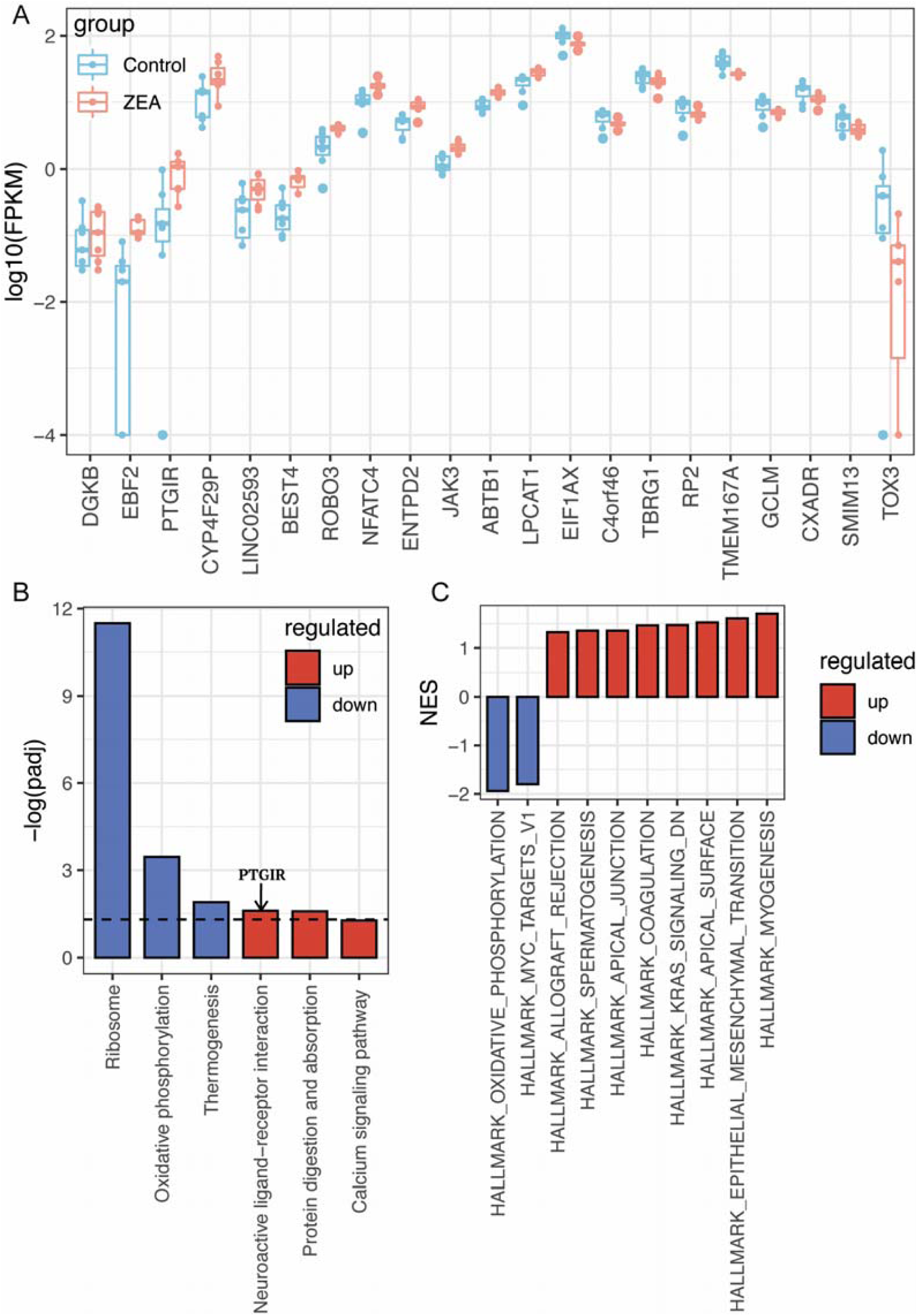
Transcriptional changes in tumor cells in response to ZEA treatment. (A) Boxplot of log-transformed expression levels (in the form of fragments per kilobase of transcript per million mapped reads [FPKM] values) of all differentially expressed genes identified by differential gene expression analysis based on the negative binomial distribution (DESeq2) at adjusted *p-value* < 0.1; (B) The histogram represents Kyoto Encyclopedia of Genes and Genomes (KEGG) pathway enrichment result for tumor sample differentially expressed mRNAs (DEmRNAs) in ZEA group compared with the control. The DE gene PTGIR was found to be involved in neuroactive ligand-receptor interaction pathways; (C) Normalized enrichment scores (NES) of the positively and negatively enriched gene sets (FDRLJ < LJ0.25) from Molecular Signatures Database (MSigDB) hallmark gene sets significantly enriched in ZEA treated mice compared to control mice.

**Table 1.** The 21 differentially expressed genes (adjusted *p-value*, a positive/negative log_2_ fold change (log_2_FC) value indicates up-regulation/down-regulation) for the comparison of the ZEA treatment versus the control, ranked by the log_2_FC.

| Gene symbol | Gene name | $\log_2$ fold change | Adjusted *p-value* |
| --- | --- | --- | --- |
| DGKB | diacylglycerol kinase beta | 4.37 | 0.039 |
| EBF2 | EBF transcription factor 2 | 2.92 | 0.054 |
| PTGIR | prostaglandin I2 receptor | 1.71 | 0.066 |
| CYP4F29P | cytochrome P450 family 4 subfamily F member 29, pseudogene | 1.37 | 0.013 |
| LINC02593 | long intergenic non-protein coding RNA 2593 | 1.36 | 0.054 |
| BEST4 | bestrophin 4 | 1.32 | 0.054 |
| ROBO3 | roundabout guidance receptor 3 | 0.84 | 0.022 |
| NFATC4 | nuclear factor of activated T cells 4 | 0.67 | 0.044 |
| ENTPD2 | ectonucleoside triphosphate diphosphohydrolase 2 | 0.63 | 0.054 |
| JAK3 | Janus kinase 3 | 0.63 | 0.045 |
| ABTB1 | ankyrin repeat and BTB domain containing 1 | 0.52 | 0.041 |
| LPCAT1 | lysophosphatidylcholine acyltransferase 1 | 0.38 | 0.005 |
| TOX3 | TOX high mobility group box family member 3 | -3.00 | 0.045 |
| SMIM13 | small integral membrane protein 13 | -0.56 | 0.018 |
| CXADR | CXADR Ig-like cell adhesion molecule | -0.46 | 0.066 |
| GCLM | glutamate-cysteine ligase modifier subunit | -0.45 | 0.027 |
| TMEM167A | transmembrane protein 167A | -0.43 | 0.013 |
| RP2 | RP2 activator of ARL3 GTPase | -0.41 | 0.068 |
| TBRG1 | transforming growth factor beta regulator 1 | -0.40 | 0.094 |
| C4orf46 | chromosome 4 open reading frame 46 | -0.39 | 0.041 |
| EIF1AX | eukaryotic translation initiation factor 1A X-linked | -0.34 | 0.025 |

We performed an enrichment analysis of the KEGG pathways and MSigDB gene sets, including (1) cancer hallmarks, (2) oncogenic signatures, and (3) chemical and genetic perturbations. The results revealed three upregulated KEGG pathways (Fig. 2B): calcium signaling (*p_adj_* = 0.055), protein digestion and absorption (*p_adj_* = 0.026), and neuroactive ligand-receptor interaction (*p_adj_* = 0.025). Downregulated pathways included ribosomes (*p_adj_* = 3.20 x 10^-12^), oxidative phosphorylation (*p_adj_*= 3.51 x 10^-4^), and thermogenesis (*p_adj_* = 0.0128) (Fig. 2B). Recent studies reported that the KEGG pathway "ribosome” (ko03010) was downregulated in colon cancer [30, 31]. Furthermore, the DE gene PTGIR is involved in the DE-upregulated neuroactive ligand-receptor interaction pathway. Regarding the MSigDB gene sets, enrichment analysis showed that ZEA intervention promoted gene signatures representing oncogenic signaling, tumor invasiveness, and metastasis, such as epithelial-mesenchymal transition (EMT), a gain of stemness, the KRAS pathway, and an invasive cancer signature (|NES score| > 1 and FDR < 0.25; Fig. 2C and Fig. S3).

### Changes in serum metabolic profiles

Both targeted and non-targeted metabolomic analyses of the serum samples were performed to explore the potential metabolic consequences of ZEA intervention. Overall, the metabolic profiles of the ZEA and control groups were separated into two clusters (Fig. 3A–B and Fig. S4A–B). Forty-one targeted metabolites were detected, of which 19 were significantly different (*p* < 0.05, independent samples t-test) between the ZEA and control groups (Fig. 3A, Fig. S4A, Table S1). The concentration of serum amino acids (AA) was lower in the ZEA intervention group than in the control group, which is consistent with two previous studies in healthy rats and porcine intestinal cell lines [32, 33]. Among the AA metabolites surveyed, phosphoethanolamine, taurine, 1-methyl histidine, and kynurenine showed the most significant changes (|log2FC| > 0.8). Additionally, 21 metabolites were identified with significant differences (*p* < 0.05, independent samples t-test) between the ZEA and control groups (Fig. 3B, Table S2). Fourteen were upregulated, including organic acids and their derivatives, indolic compounds, purine derivatives, and ethanolamine, whereas the remaining were downregulated, mainly lipids and quinazoline. Among the identified metabolites, coumarinic acid, uric acid, and malic acid showed the largest fold changes (|log_2_FC| > 1).

**Fig. 3.**
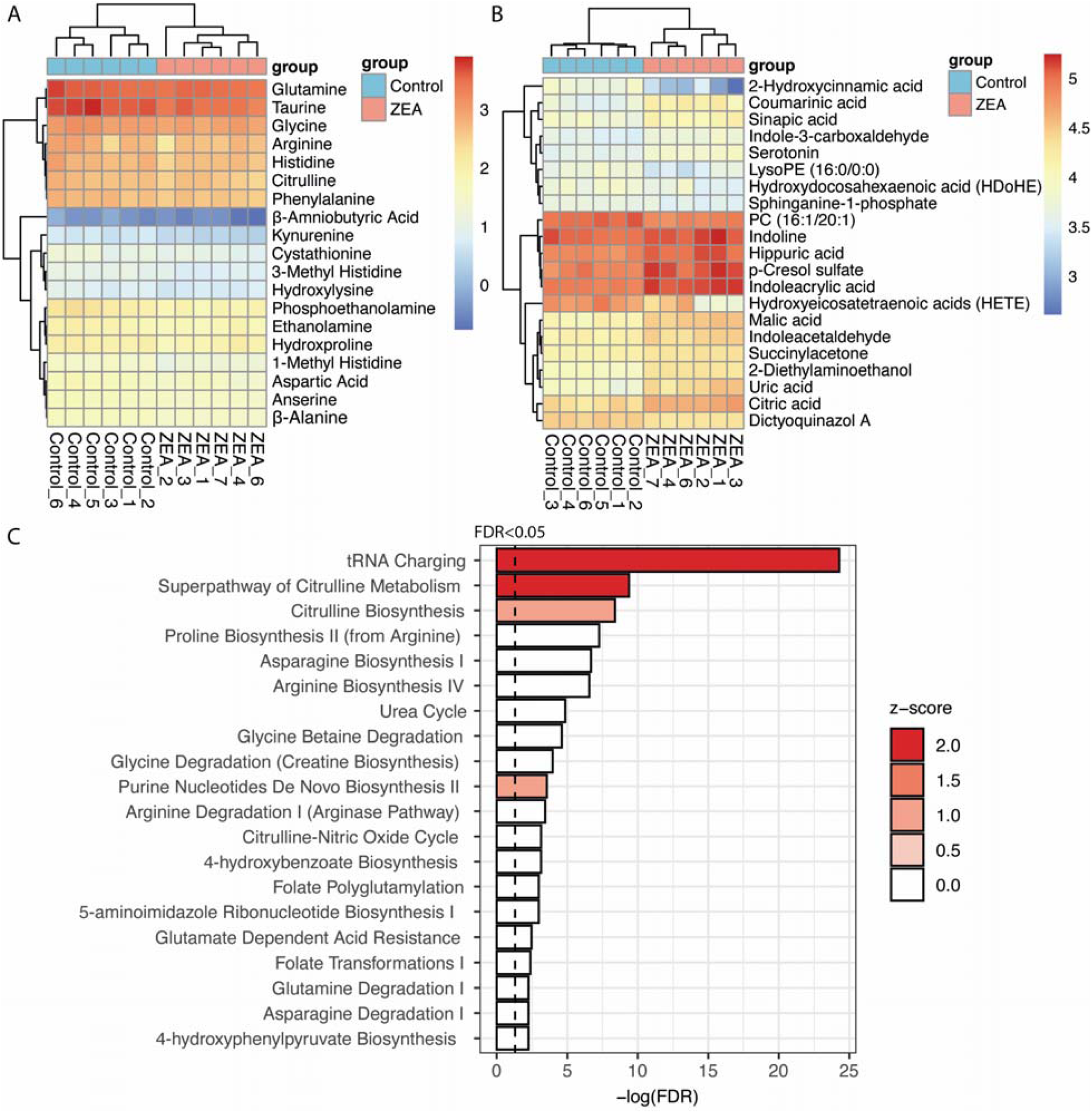
Effect of ZEA intervention on metabolites in serum samples. (A) Heatmap with log-transformed levels of 19 differentially abundant metabolites between the ZEA and control groups following targeted metabolomics analysis. (B) Heatmap with log-transformed expression levels of 21 identified differentially abundant metabolites between the ZEA and control groups following non-targeted metabolomics profiling. (C) Top 20 canonical pathways identified with the -log(FDR) and z-score listed. The threshold was set at 1.3 for -log(FDR) (equivalent to FDR < 0.05).

To investigate significantly altered metabolic pathways, the log2FC of each metabolite was calculated using Ingenuity Pathway Analysis for canonical pathway analysis. The top 20 canonical pathways were identified and ranked according to their false discovery rates (FDRs) (Bonferroni corrected *p-values*) (Fig. 3C). Among these pathways, four activated pathways, including tRNA charging, the super pathway of citrulline metabolism, citrulline biosynthesis, and the purine nucleotide de novo biosynthesis II pathway, were identified with positive z-scores in the ZEA treatment group compared to the control group. The enriched citrulline metabolism and biosynthesis pathways indicated that ZEA intervention influenced amino acid metabolism in mice, consistent with a previous study using a breast cancer cell line [34].

### Changes in gut microbiota and microbiota-derived SCFAs

The gut microbiota’s alpha diversity (Shannon index) was estimated based on the taxonomic profiles at the OTU level. The results indicated a significantly lower (*p* = 0.007 with an independent-sample t-test) alpha diversity in the ZEA treatment group than in the control group (Fig. 4A). Moreover, the MDS ordinations based on beta diversity (robust Aitchison distance) showed a difference between the two groups to a certain extent (R = 0.1254, *p* = 0.096 with ANOSIM) (Fig. 4B). The Aitchison distances between samples in the ZEA-treated group were significantly larger than those in the control group (*p* = 0.006, Wilcoxon test), suggesting that the microbial communities became more exhibited greater divergence following ZEA intervention. Nine genera were significantly differentially abundant (DA) between the ZEA-treated and control groups, based on ANCOM-BC (Fig. 4C). The relative abundance of *Alistipes, Candidatus Saccharimonas, Helicobacter, Lachnospiraceae UCG-010, Paludicola, Rikenella, Rikenellaceae RC9 gut group,* and *Tuzzerella* was significantly lower in the ZEA treatment group than in the control group (Fig. 4C, Fig. S5). *Rikenella* is an SCFA producer among the differentially abundant genera, and *Candidatus Saccharimonas* is associated with anti-inflammation [35]. Together, these results suggest that ZEA contributes to tumor growth by repressing SCFA-producing or anti-inflammatory bacteria.

**Fig. 4.**
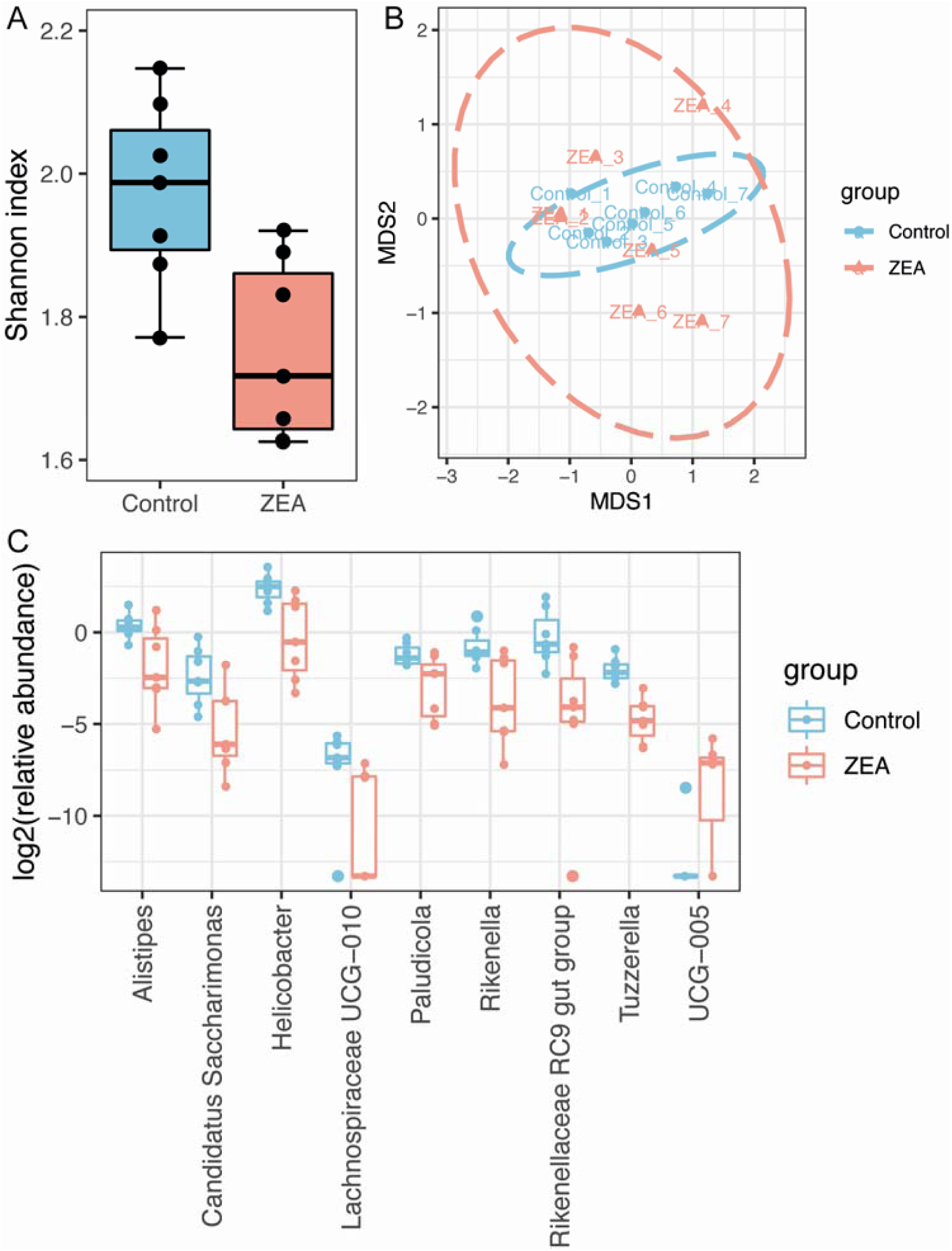
Altered cecal microbiome community upon ZEA treatment. (A) Boxplot of alpha-diversity indices. Shannon index represent the community diversity of the microbiota in the cecal content. Boxes represent the interquartile range (IQR) between the first and third quartiles (25th and 75th percentiles, respectively), and the horizontal line inside the box defines the median. ** *p* < 0.01 (*p* = 0.007, independent-sample t-test). (B) Multiple dimension scale (MDS) plots showing the beta diversity results of cecal microbial profiles between the control and ZEA groups based on robust Aitchison distance. (ANOSIM statistic R 0.1254, *p* = 0.096, based on 999 permutations). (C) Boxplot of log-transformed relative abundance of significantly (FDR < 0.05) different genera between the control and ZEA detected by ANCOM-BC.

We further assessed the effect of ZEA treatment on major gut microbiota-derived SCFAs (acetic acid, propionic acid, isobutyric acid, butyric acid, and valeric acid) using GC-MS. ZEA treatment reduced SCFA production compared to the control group (Fig. S6). The concentrations of total SCFAs, acetic acid, propionic acid, and butyric acid in the ZEA treatment group were significantly lower (*p* < 0.05, independent-sample t-test) than those in the control group (Fig. S6).

### Associations among gene expression, metabolites, and gut microbes

Thirteen metabolites were significantly correlated with tumor weight using Spearman’s correlation (Table S3). Among them, six identified metabolites, including indolic compounds and their derivatives (serotonin and indole acetaldehyde), purine derivatives (uric acid), and organic acid derivatives (p-cresol sulfate, succinylacetone, and hippuric acid), were positively correlated with tumor weight (correlation > 0.6, *p* < 0.05). In contrast, other metabolites, including two amino acids (1-methyl histidine and β-alanine) and lipid-related molecules hydroxydocosahexaenoic acid (HDoHE) and hydroxyeicosatetraenoic acids (HETE), were negatively correlated (correlation < -0.6, *p* < 0.05) with tumor weight.

Spearman’s correlation coefficients were calculated to establish links among differentially expressed genes, differentially abundant metabolites, and differentially abundant gut microbes. A total of 12 significantly and strongly positive (FDR < 0.05, |correlation|> 0.8) and 21 significantly and strongly negative correlations were identified in the integration analysis (Fig. 5A, Table S4). These 33 associations comprised of 36 features (12 differentially expressed genes, 19 differentially abundant metabolites, and five differentially abundant genera) (Fig. 5A and B). The majority (25 of 33, or 75.8%) of the associations were between metabolites and gene expression. Among the 19 differentially abundant metabolites, ten were significantly correlated with tumor weight. More specifically, indolic compounds and derivatives (serotonin and indole acetaldehyde), purine derivatives (uric acid), and organic acid derivatives (p-cresol sulfate, succinylacetone, hippuric acid, and 2-hydroxycinnamic acid) were positively correlated with tumor weight. In contrast, two amino acids (1-methyl histidine and β-alanine) and one lipid-related molecule (HETE) were negatively correlated with tumor weight. Our results revealed a strong connection between host metabolism and the metabolic potential of gut microbes.

**Fig. 5.**
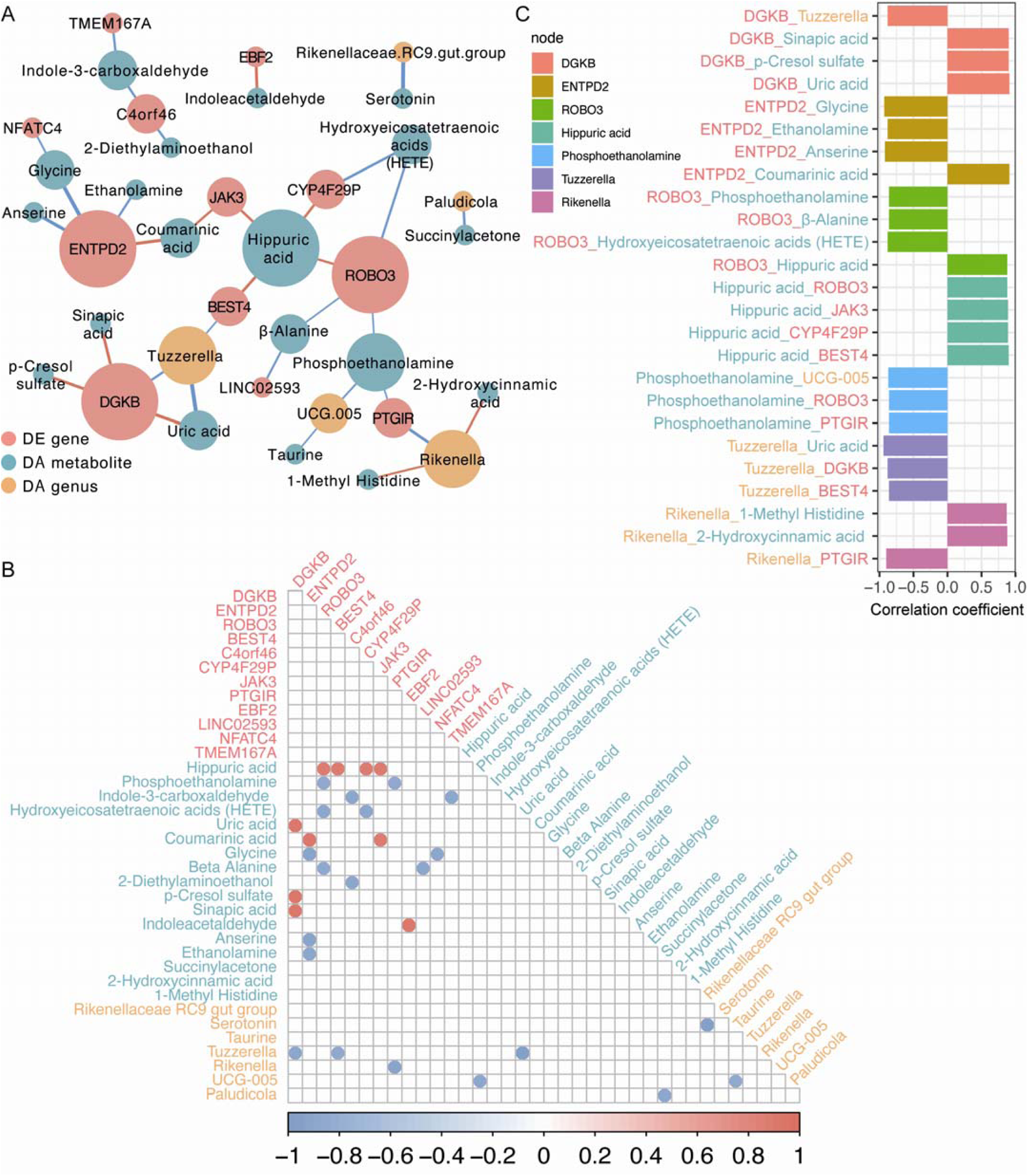
Integrative analysis of the differentially expressed genes, differentially abundant genera, and differentially abundant metabolites using Pearson correlation. (A) Association network showing the Pearson correlation analysis of differentially expressed genes (red nodes), differentially abundant genera (orange nodes), and differentially abundant metabolites (dark green nodes). The significant correlations (Bonferroni corrected FDR < 0.05) are shown in the network. Red lines represent positive correlations, and blue lines represent negative correlations; (B) Corrplot depicting the correlations among differentially expressed genes, differentially abundant genera, and differentially abundant metabolites. Only features showing the correlations of > 0.8 or < −0.8 in any of the pairwise associations were included for visualization. The significance level was set at an FDR < 0.05; (C) Boxplot showing correlation of seven nodes whose degree ≥ 3 in Fig. 5(A). DGKB, diacylglycerol kinase beta; ENTPD2, ectonucleoside triphosphate diphosphohydrolase 2; ROBO3, roundabout guidance receptor 3; Hippuric acid; Phosphoethanolamine; *Tuzzerella; Rikenella*.

Seven nodes had at least three associations with other features (Fig. 5A and C). Among these nodes, the differentially expressed gene DGKB was strongly (|correlation| > 0.8) positively correlated with uric acid (correlation = 0.909), p-Cresol sulfate (correlation = 0.902) and sinapic acid (correlation = 0.902), but negatively correlated with *Tuzzerella* (correlation = -0.881) (Fig. 5 A-C), indicating that the interplay among host metabolites, gut microbiota and tumor gene expression may contribute to the ZEA-induced tumor growth. Among the metabolite features, hippuric acid was frequently associated with multiple differentially expressed genes with oncogenic potential (positive correlation with BEST4 [correlation = 0.895], CYP4F29P [correlation = 0.888], JAK3 [correlation = 0.888], and ROBO3 [correlation = 0.881]) (Fig. 5A– C). *Tuzzerella* showed the highest number of associations with other genes and metabolites. *Tuzzerella* was negatively associated with BEST4 (correlation = -0.86), DGKB (correlation = - 0.881), and uric acid (correlation = -0.937) (Fig. 5 A-C). The identified gene-metabolite-genus crosslinks provide mechanistic insights into zearalenone-induced tumor growth in CRC.

## Discussion

To the best of our knowledge, this is the first study to provide evidence that ZEA promotes CRC growth *in vivo*. Through comprehensive and integrative multi-omics analysis, we revealed that ZEA promotes CRC tumor growth through multiple processes, including promoting the BEST4/AKT/ERK1/2 pathway, modifying serum metabolites, and modulating gut microbiota composition.

To explore the influence of ZEA on colon tumor growth, we performed RNA-seq to identify the DEGs. We identified two ZEA-promoting genes (BEST4 and DGKB), which were previously reported to be related to the malignant growth of colon cancer. RTLJPCR analysis confirmed that BEST4 was significantly up-regulated in the ZEA group (*p* = 0.024, independent samples t-test; Fig. S7). IHC staining revealed significantly increased levels of BEST4, pERK1/2, pAKT, and DGBK in the ZEA group (Fig. S8A-B), which agrees with our previous studies [10]. BEST4 overexpression is correlated with a more advanced TNM stage, lymph node metastasis, and a lower survival rate. Through activating the BEST4/PI3K/AKT signaling pathway, BEST4 promotes CRC proliferation and migration [29].

We conducted non-targeted and targeted metabolomic analyses of serum samples to reveal the metabolic influence of ZEA exposure. Consistent with previous findings, which revealed that ZEA alters amino acid metabolism [32, 33], the serum amino acid (AA) concentration was lower in ZEA-treated group (Table S1-2). However, the mechanism through which ZEA modulates amino acid metabolism remains unclear. One potential explanation could be the higher metabolic demand in the ZEA group owing to the high tumor weight, supported by the enriched KEGG pathway and protein digestion and absorption. Correlation analysis was performed to identify altered metabolites that could contribute to transcriptomic changes in the ZEA group. DGKB is positively correlated with p-cresol sulfate, which has been previously reported to induce inflammation and tumor growth in CRC [36]. P-cresol sulfate is derived from microbial metabolism and conjugation of AAs, phenylalanine, and tyrosine in the colon [37]. This finding could explain the significantly lower phenylalanine levels in the ZEA group because phenylalanine could be metabolized or converted to tyrosine following microbial conversion. In addition to amino acids, the concentrations of lipid-related metabolites were lower in the ZEA group (Fig. 3B) and negatively correlated with tumor weight (Table S3). We observed that serum HETE level was lower in the ZEA group, potentially leading to defects in peroxisome proliferator-activated receptor γ activation, which is responsible for apoptosis in CRC [38]. Another lipid-related metabolite, HDoHE, was also negatively correlated with tumor weight, in line with a previous study that revealed a higher level of HDoHE was associated with smaller CRC tumors in mice, owing to the anti-inflammatory properties of HDoHE [39]. These findings indicate that ZEA may aggravate tumor growth by interfering with systemic metabolism.

Alteration in the microbial community are known to influence the development of diseases such as irritable bowel disease, obesity, liver disease and cancer [40–42]. Consistent with a previous study that revealed a significantly lower cecal SCFAs content in the ZEA-treated animal group [43], we observed reduced cecal SCFAs and SCFA-producing bacteria, *Rikenella* [44] and *Rikenellaceae RC9 gut group*, and the anti-inflammatory bacterium *Candidatus Saccharimonas* [35]. A low abundance of SCFAs, major metabolites from the gut microbiota, and SCFA-producing bacteria has been linked to colon cancer development [45]. Additionally, our correlation analysis revealed that the abundance of *Tuzzerella*, which was previously reported to activate the PI3K/AKT pathway, was negatively correlated with serum uric acid levels [46]. The metabolomic role of *Tuzzerella* and its contribution to in tumor pathogenesis remain unclear. Further studies are needed to elucidate its metabolic influence and potential interplay with the expression of BEST4.

We showed that ZEA promoted tumor development by altering the gut microbiota, microbiota-derived metabolites, and oncogenic gene expression during CRC progression. However, this study had several limitations. One major limitation of this study was that the animal models used may not provide the same human responses upon exposure to ZEA. Although humans and mice share similar physiological and anatomical structures, only 15% of gut commensal bacteria are shared. Most human microbiome studies have used fecal samples (as human ferment indigestible food components in the colon), whereas the cecum is the site of fermentation in mice [47].This may lead to discrepancies when translated to clinical settings.

Additionally, we were unable to distinguish between the bacterial species and identify the metabolic features of the microbiome owing to the limitations of 16S rDNA sequencing. To estimate the effect of ZEA on human subjects with CRC more accurately, we suggest a humanized mouse model with shotgun metagenomic sequencing.

## Conclusions

In summary, this study, for the first time, provided mechanistic insights into how ZEA promotes tumor development in colon cancer. We demonstrated that exposure to ZEA, even at a low dose, substantially altered the tumor transcriptome, serum metabolome, and gut microbiota. ZEA-induced tumor progression is likely mediated by modulation of the BEST4/AKT/ERK1/2 pathway, circulatory amino acid levels reduction, and alteration of gut microbiota composition. Through the integration and correlation analysis of multi-omics data, we found that *Tuzzerella* was significantly negatively correlated with BEST4 expression levels, and the serum metabolite

uric acid was positively correlated with tumor weight. Since ZEA is commonly found in the human diet and its impact can be persistent, further studies on the potential contribution of chronic exposure to ZEA to colon cancer development are warranted.

## Compliance with Ethics Requirement

All the Institutional Guidelines for the care and use of animals were followed. Animal handling was approved by the Committee on the Use of Live Animals in Teaching and Research (CULATR No. 4785-18) at the University of Hong Kong.

## Authors’ contributions

Conceptualization, E.K.K.L. and H.E.; methodology, E.K.K.L., X.W.W, P.K.L., H.W.; investigation, E.K.K.L., X.W.W., P.K.L., H.C.W., D.Z., J.L. and H.E.; writing - original draft, E.K.K.L., X.W.W, writing - review and editing E.K.K.L., X.W.W., P.K.L, J.C.Y.L, C.G.G., D.Z, J.L. and H.E.; funding acquisition, D.Z, J.L. and H.E. All authors have read and agreed to the published version of the manuscript.

## Acknowledgements

The authors thank Mr. Oscar Chow for providing technical assistance. The authors are grateful to Dr. Sam Leung Kin Sum for providing the preliminary data on SCFAs. Fig. 1A was created with BioRender.com. This study was supported by the Shenzhen Basic Research Program (JCYJ20190808182402941), Guangdong Basic and Applied Research Major Program (2019B030302005), Collaborative Research Fund (C7013-19GF) in Hong Kong, City University of Hong Kong Internal Grant (7005756, 9678247, 9680310), and PolyU Start-up Fund (1-BE3H).

## Supplementary material

Supplementary data for this article can be found at https://doi.org/xxxx.

**Fig. S1:**
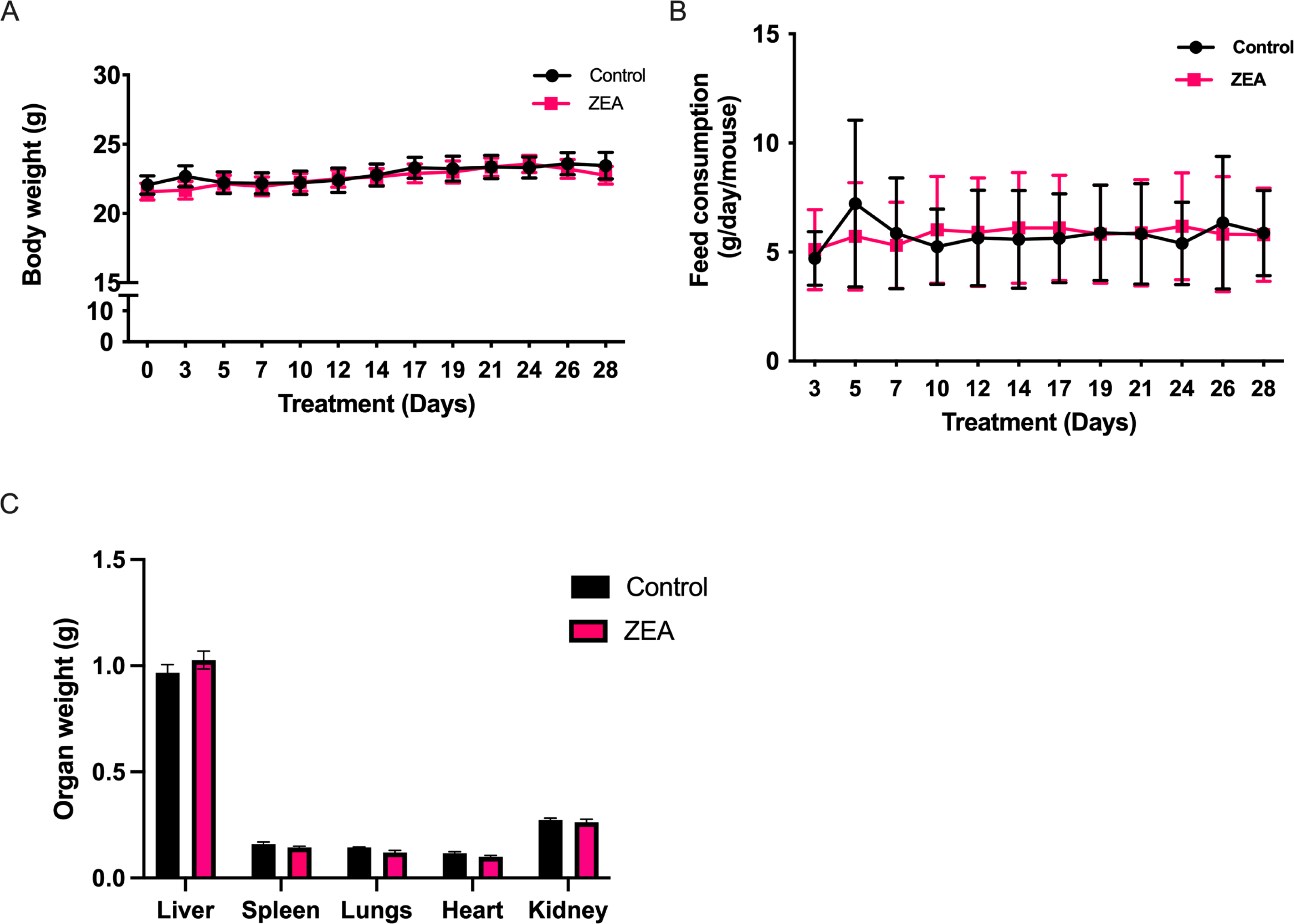
Effect of Zearalenone in body weight, feed consumption and vital organ weight of xenograft mice

**Fig. S2:**
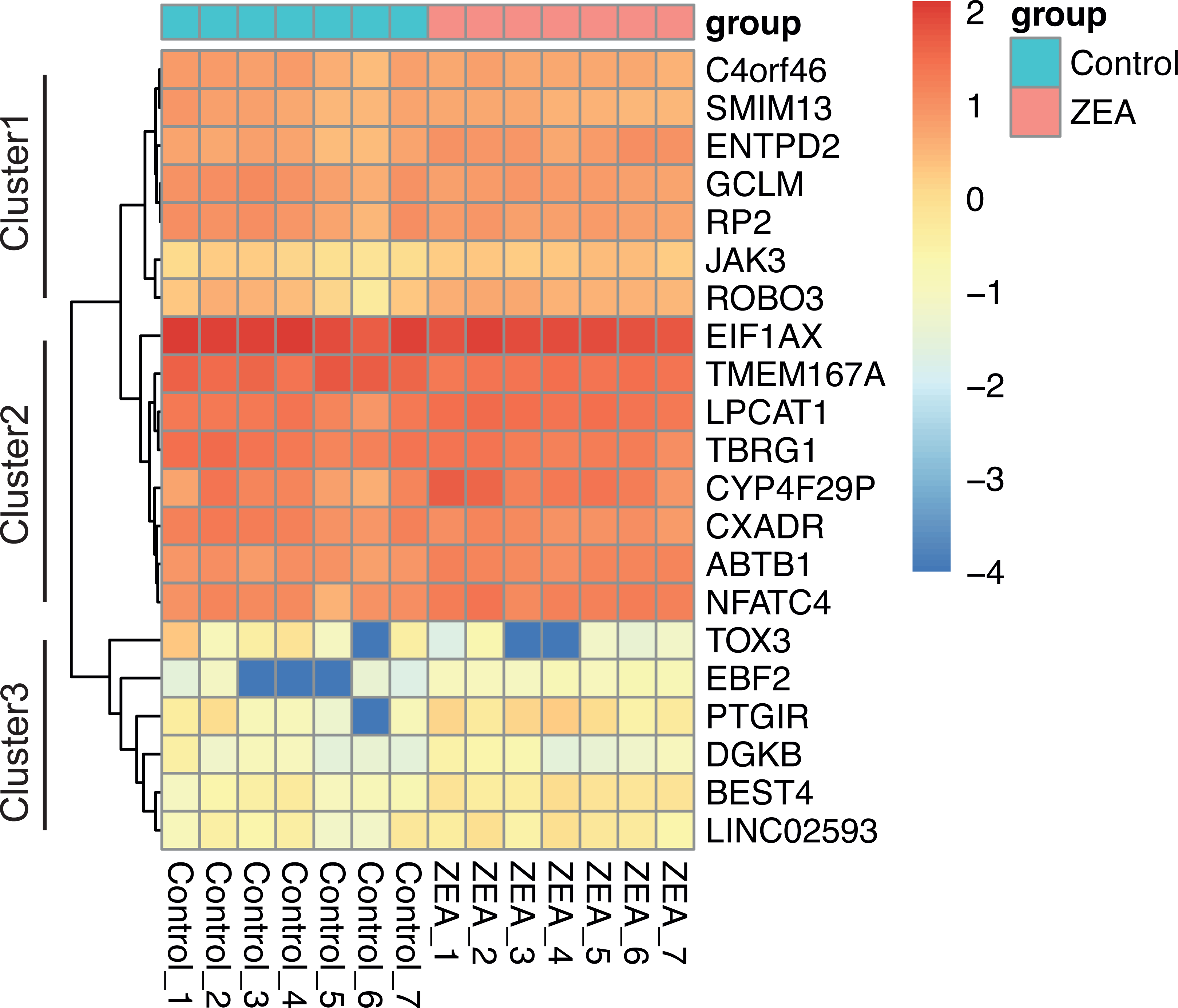
Heatmap of log-transformed expression levels (in the form of FPKM values) of all differentially expressed genes identified by DESeq2 at *padj* < 0.1

**Fig. S3:**
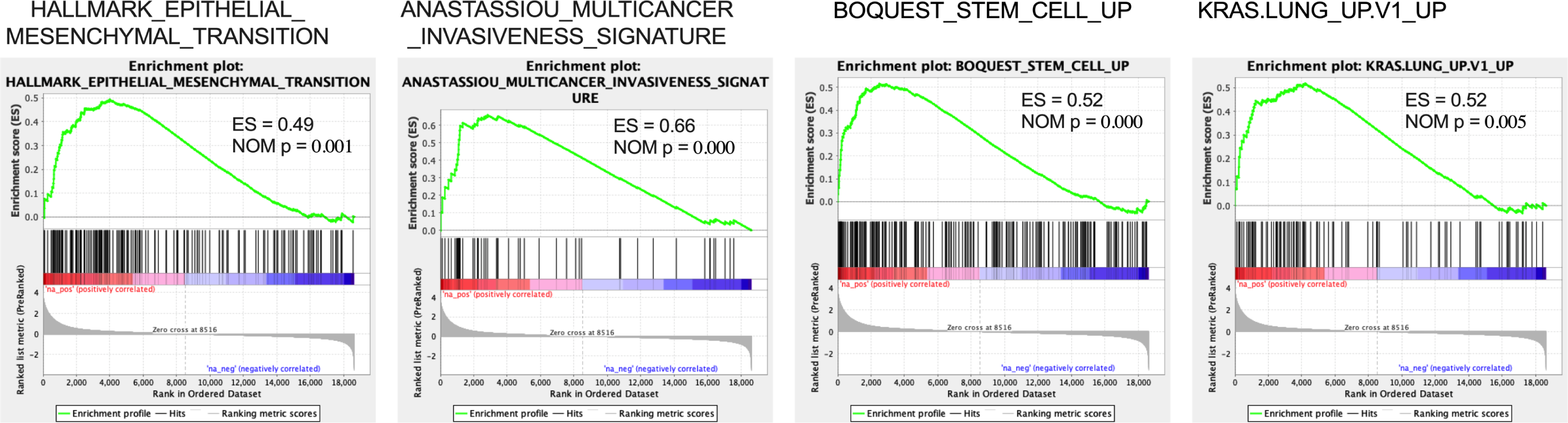
Correlations between ZEA treatment and potential oncogenic signaling

**Fig. S4:**
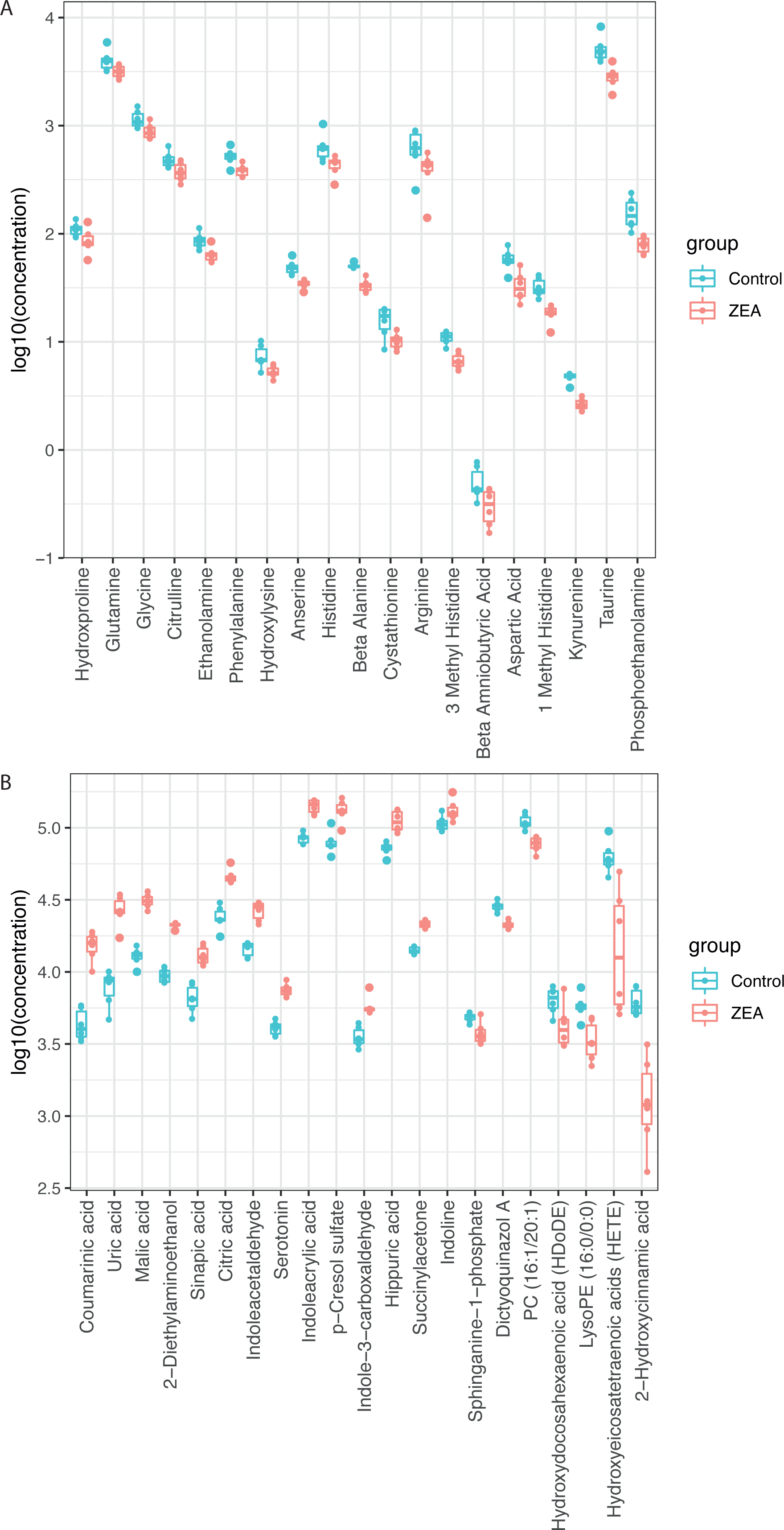
Effect of ZEA treatment on metabolites in serum samples

**Fig. S5:**
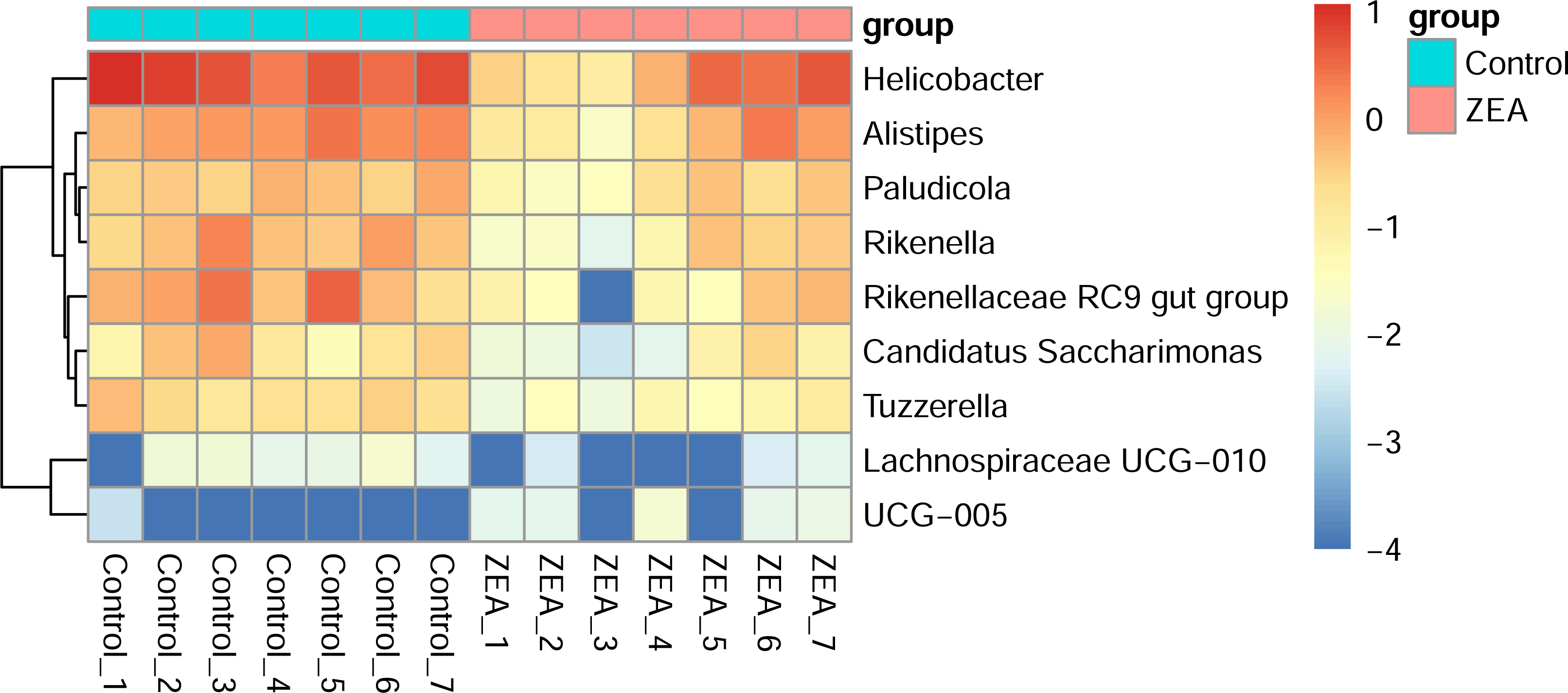
Heatmap of log-transformed relative abundance of significantly (FDR < 0.05) different genera between control and ZEA detected by ANCOM-BC

**Fig. S6:**
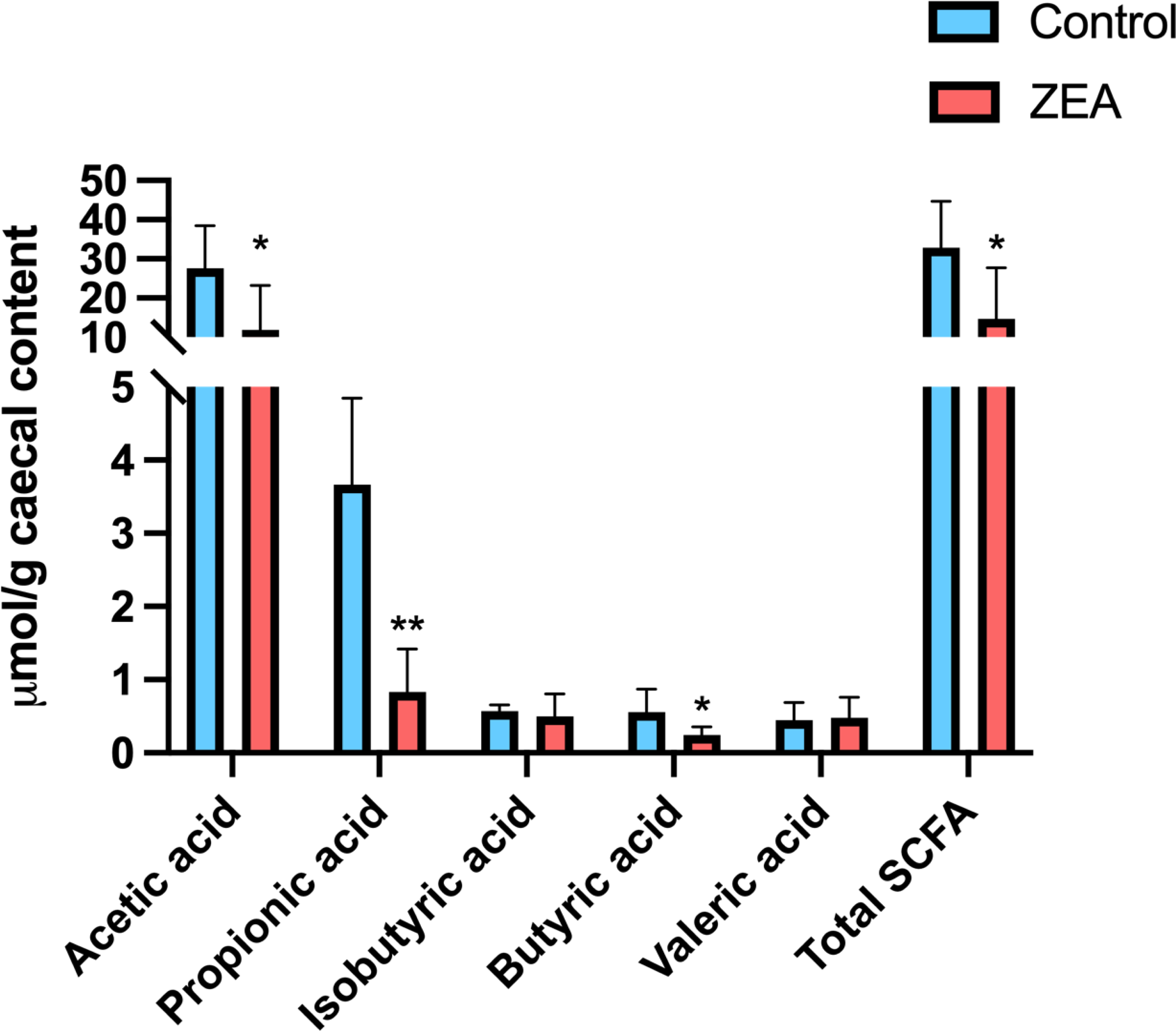
Effect on the SCFA levels in the cecal contents upon treatment

**Fig. S7:**
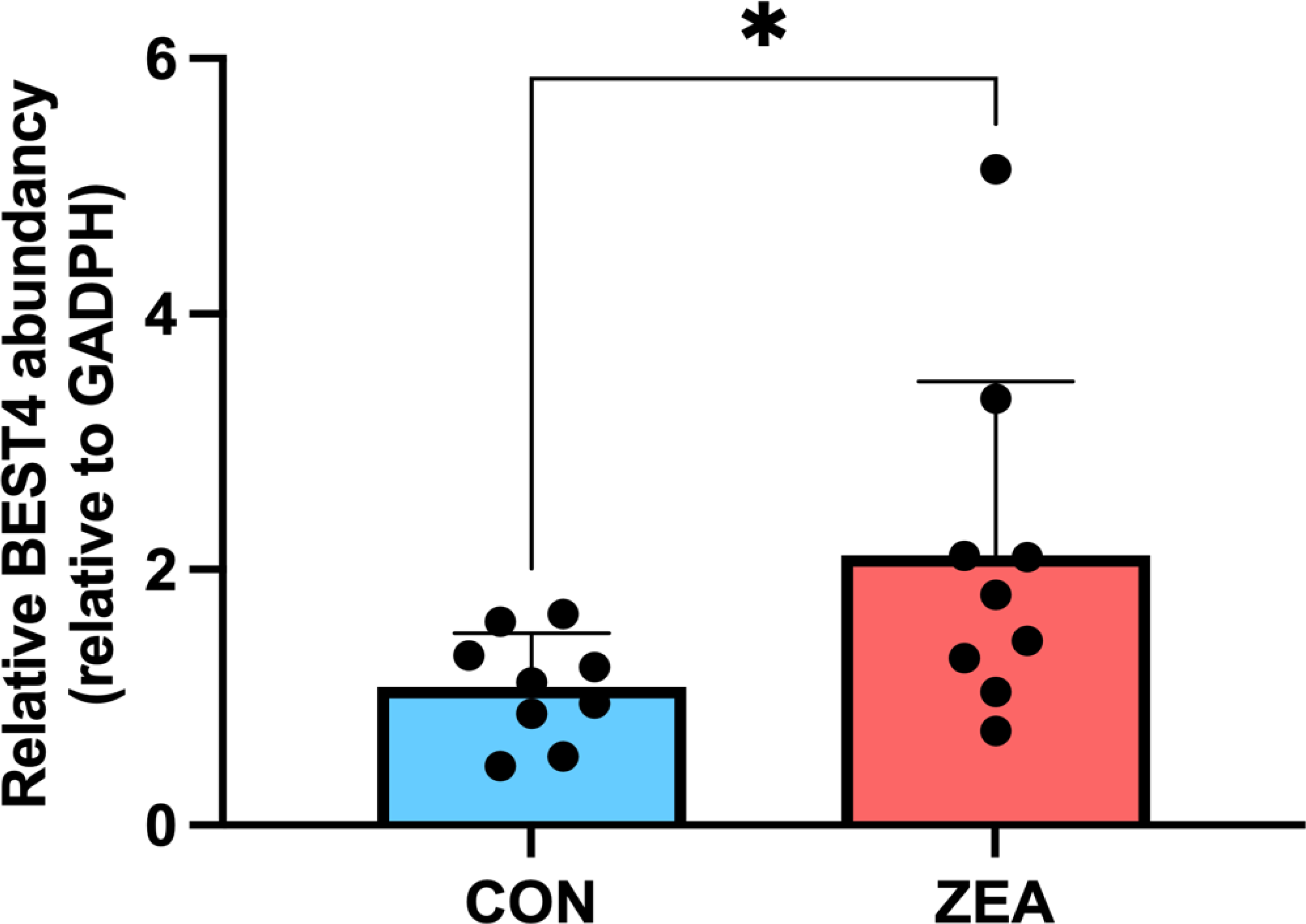
qPCR analysis of the mRNA expression of BEST4

**Fig. S8:**
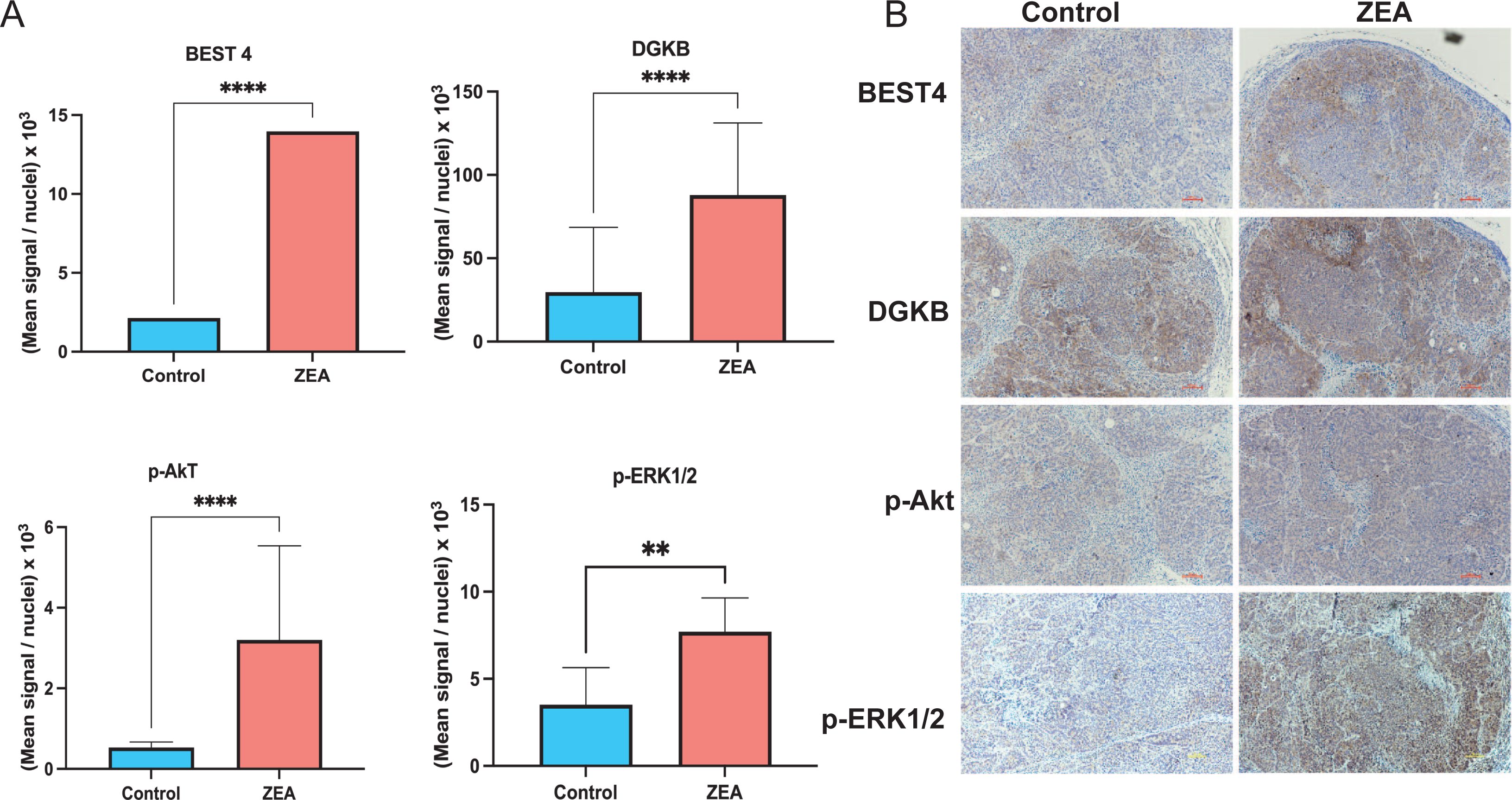
Transcriptional changes in tumor cells in response to ZEA treatment

Table S1: Amino acids that were altered by ZEA treatment compared to control group; Table S2: List of metabolites detected in serum samples of control and experimental groups using Progenesis QI; Table S3: Metabolites strongly related to tumor weight with spearman’s rank correlation coefficient; Table S4: Pearson correlation coefficient and significance level for DA metabolites, DE genes and DA genera; Supplementary methods: RNA extraction and Real time reverse transcription polymerase chain reaction (RT-qPCR).

