## supplementary materials for "Mechanistic insights into zearalenone-accelerated colorectal cancer in mice using integrative multi-omics approaches"

Supplementary Results

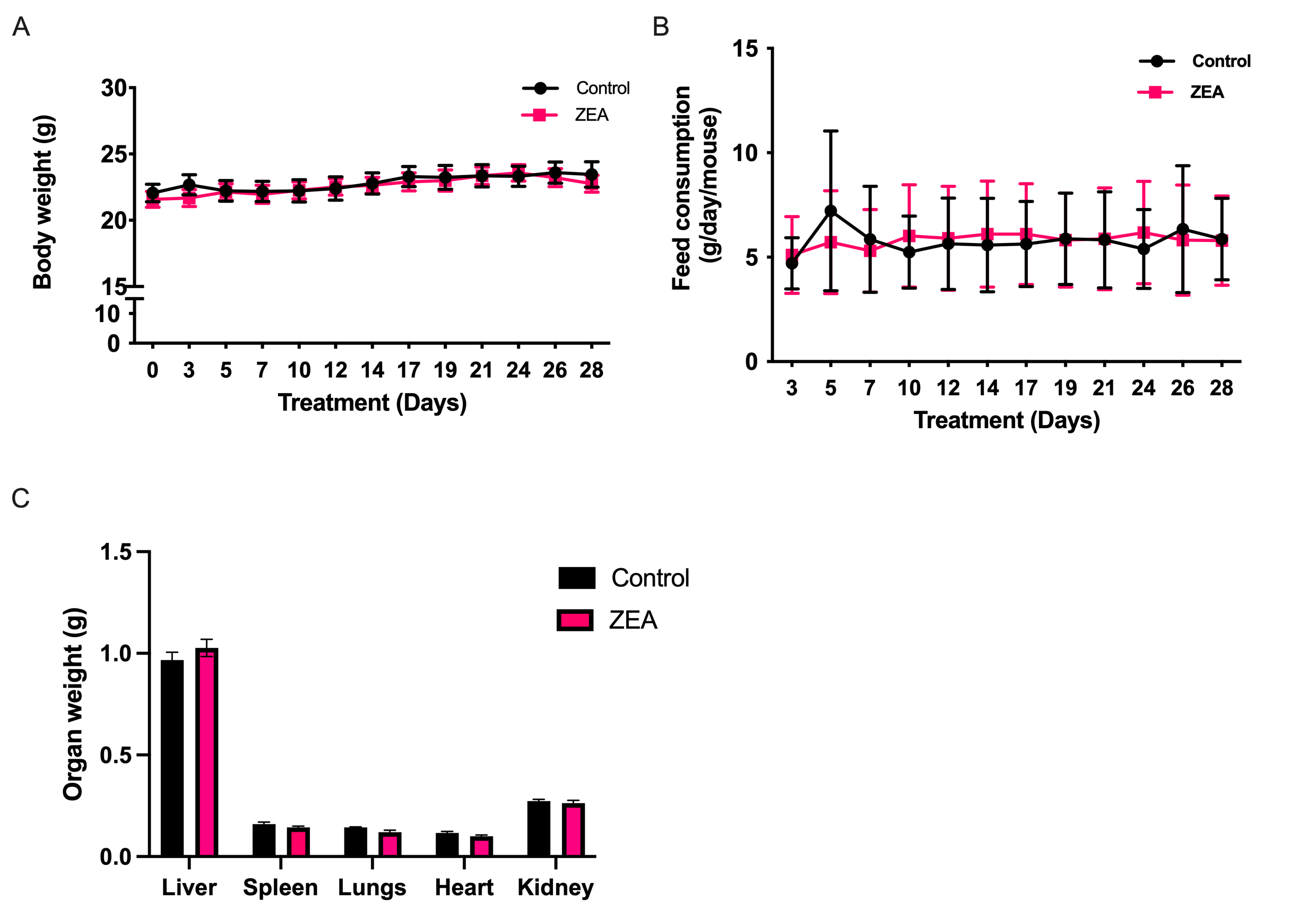

**Fig. S1**. Effect of Zearalenone in body weight, feed consumption and vital organ weight of xenograft mice. No significant changes in body weight (A), feed intake (B). Results shown are mean ± SEM, n=9. No Significant changes in organ weight of vital organs (C) including liver, spleen, lungs, heart and kidney were found (n=3).

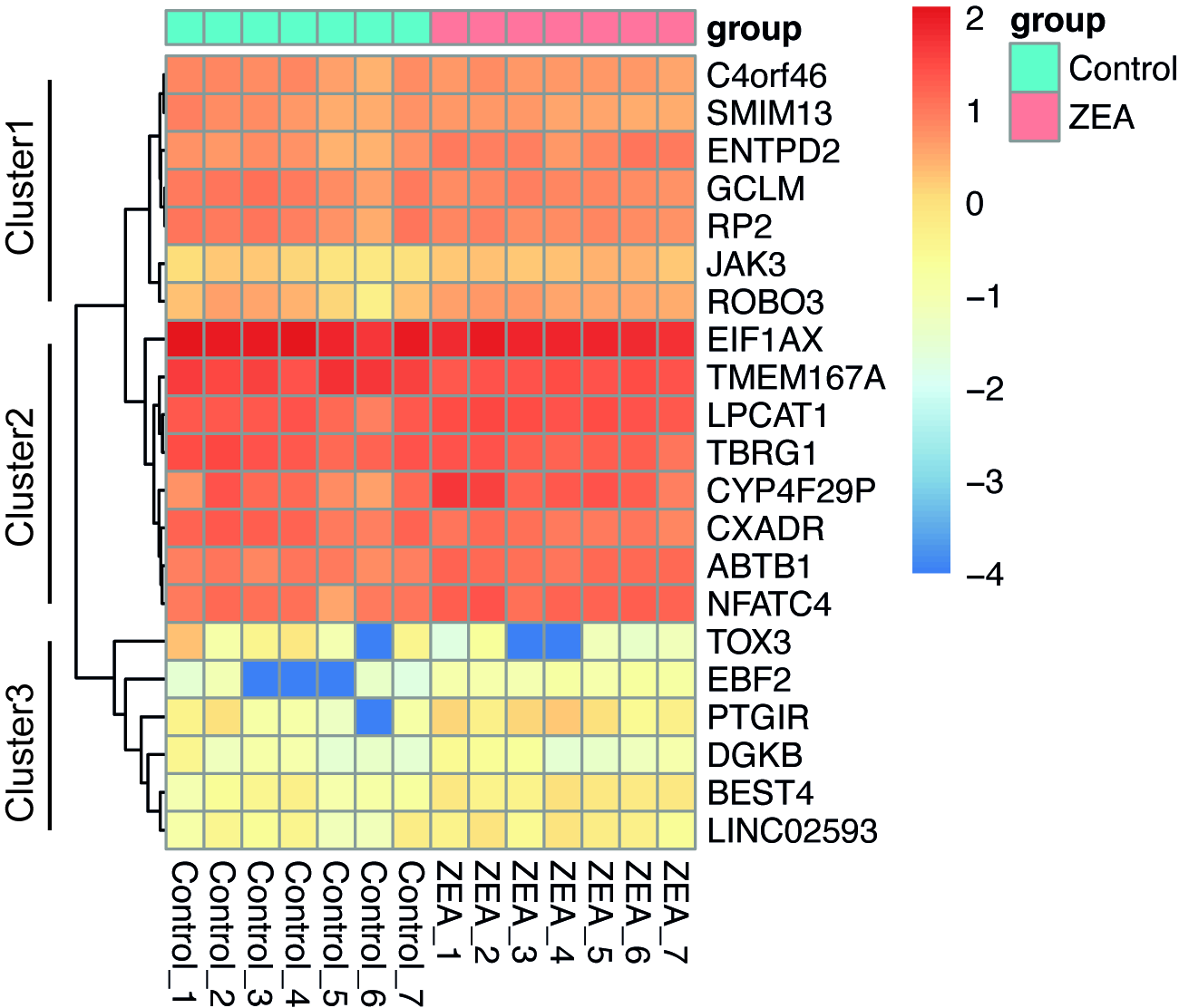

**Fig. S2.** Heatmap of log-transformed expression levels (in the form of FPKM values) of all differentially expressed genes identified by DESeq2 at padj <0.1.

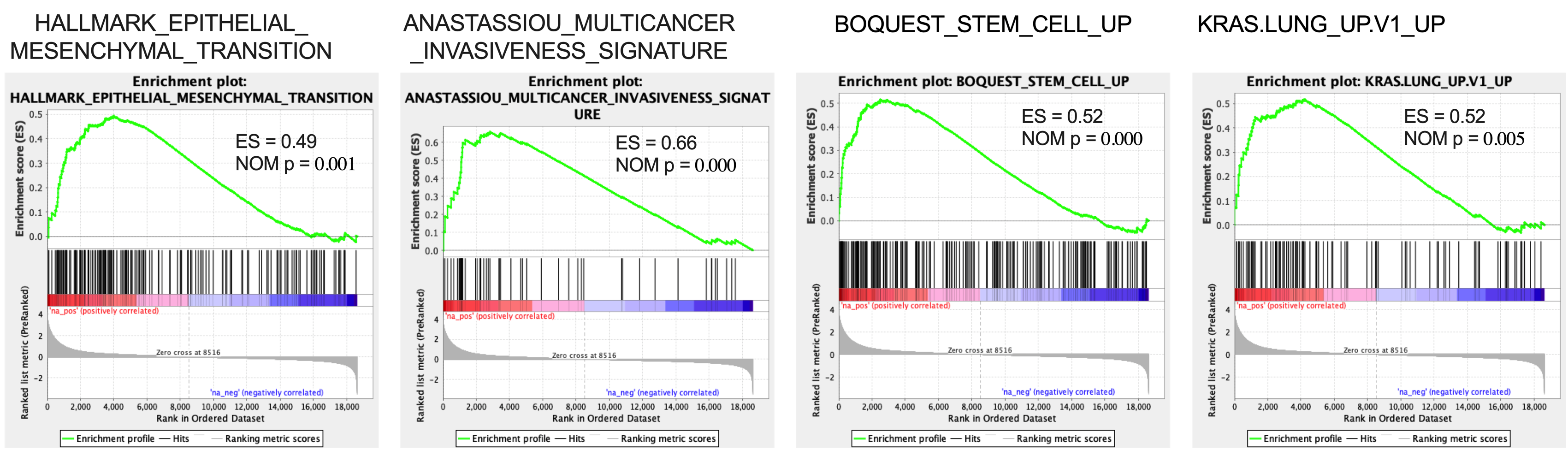

**Fig. S3.** Correlations between ZEA treatment and potential oncogenic signaling. Gene signatures indicating the epithelial mesenchymal transition, cancer multicancer insaneness, stemness and KRAS signaling pathway were significantly enriched in ZEA group compared to control, as shown by gene set enrichment analysis (GSEA).

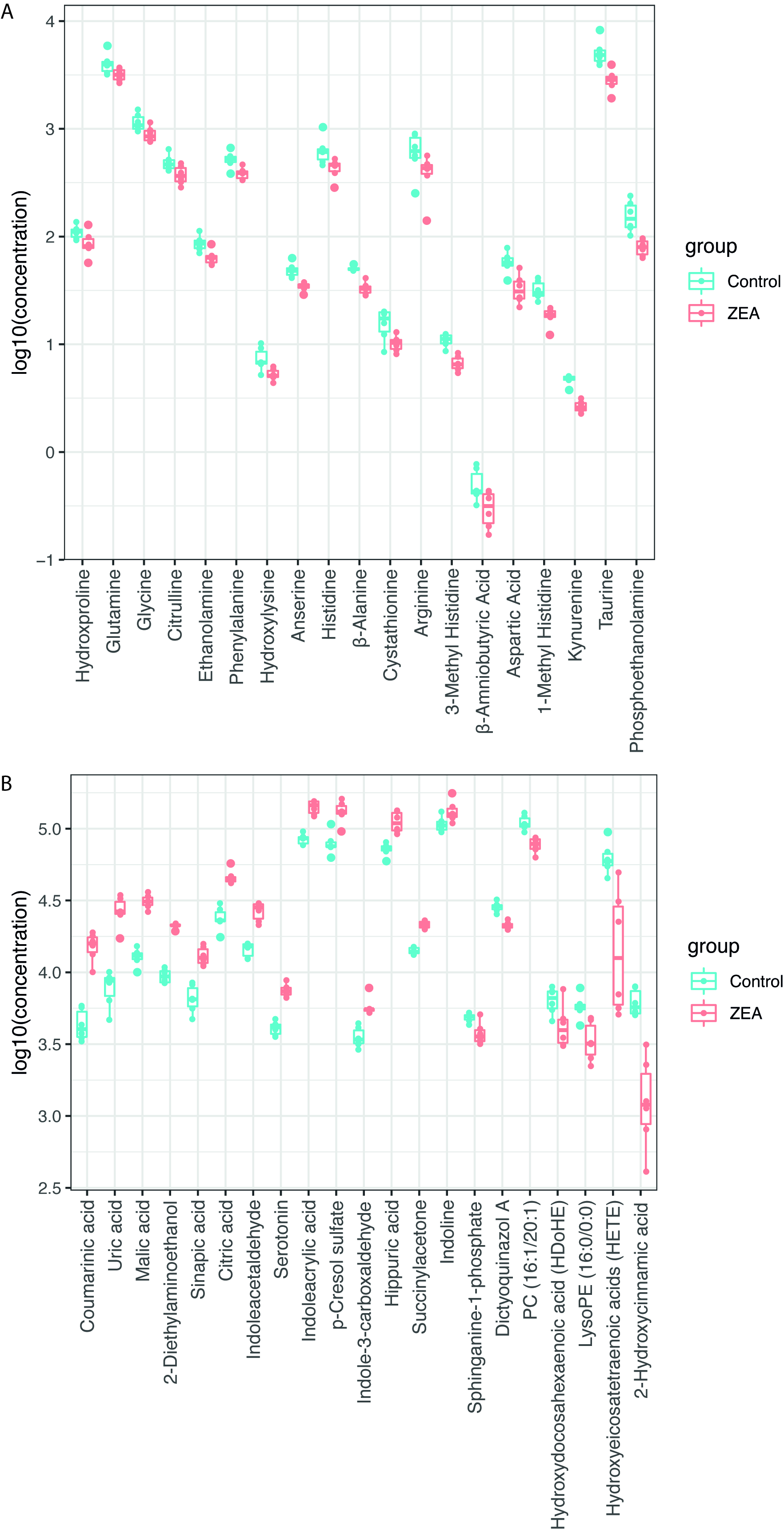

**Fig. S4.** Effect of ZEA treatment on metabolites in serum samples. (A) Boxplot with log-transformed expression levels of 19 identified differential targeted metabolites between the ZEA treatment and control groups. (B) Boxplot with log-transformed expression levels of 21 identified differential non-targeted metabolites between control and ZEA.

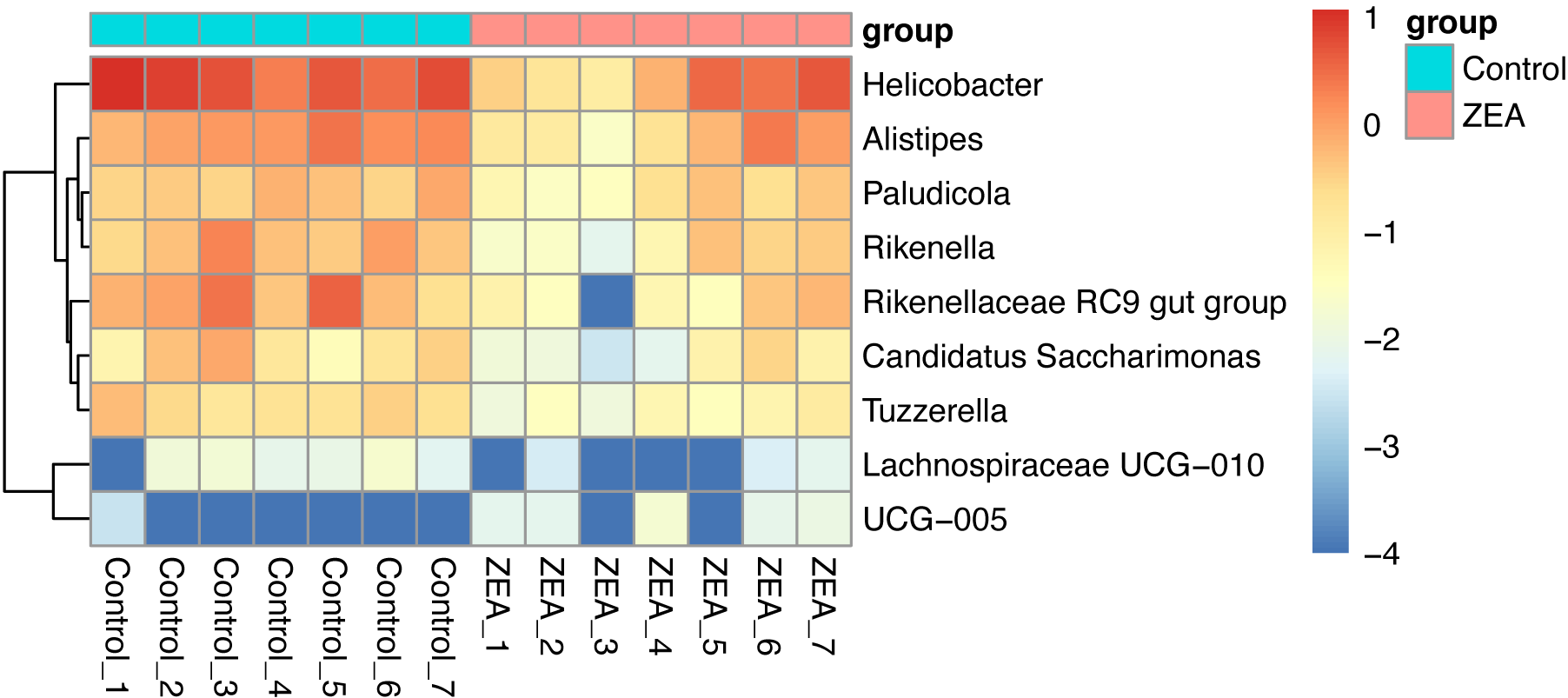

**Fig. S5.** Heatmap of log-transformed relative abundance of significantly (FDR <0.05) different genera between control and ZEA detected by ANCOM-BC.

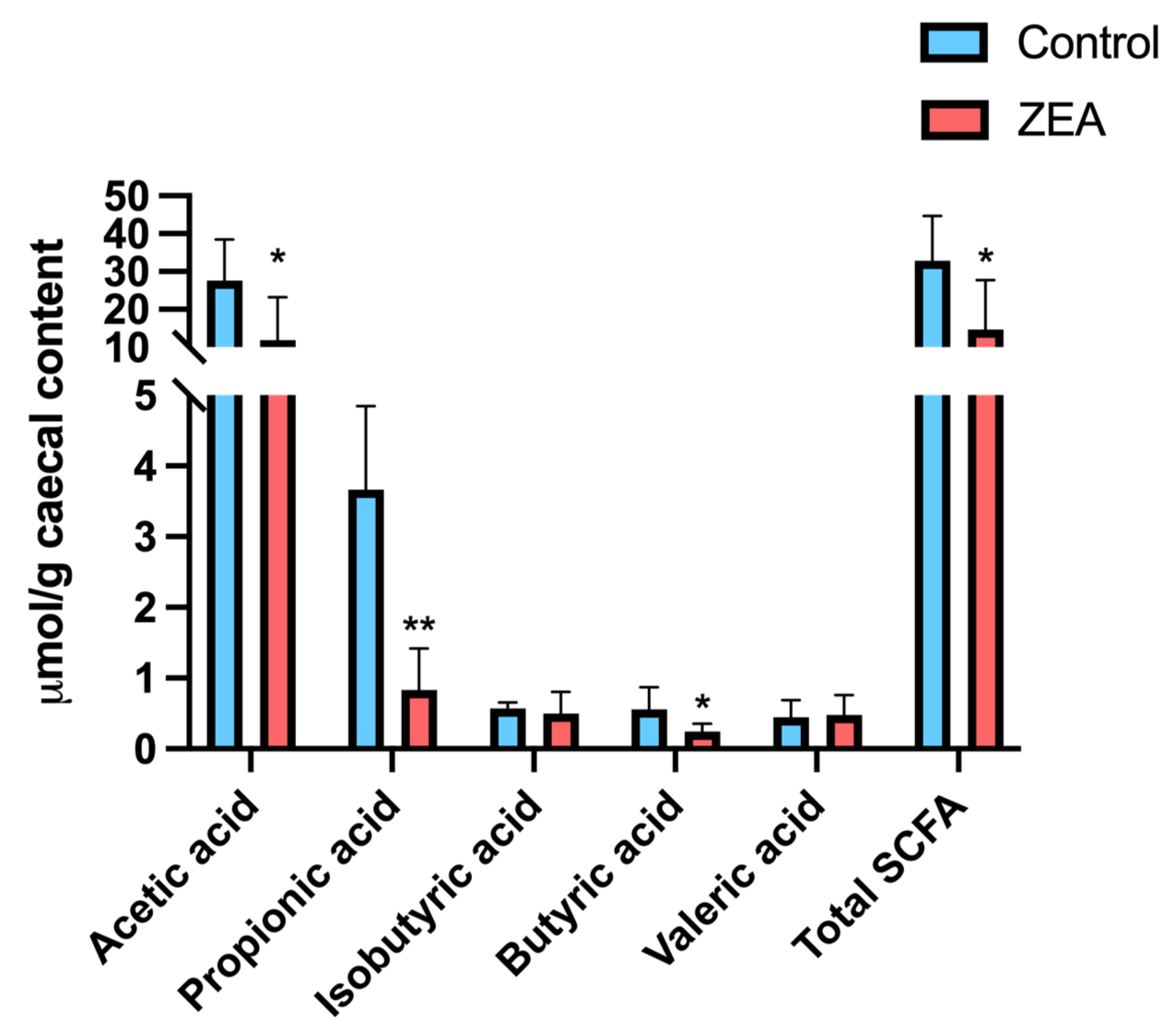

**Fig. S6.** Effect on the SCFA levels in the caecal contents upon treatment. Results shown are mean ± SEM.

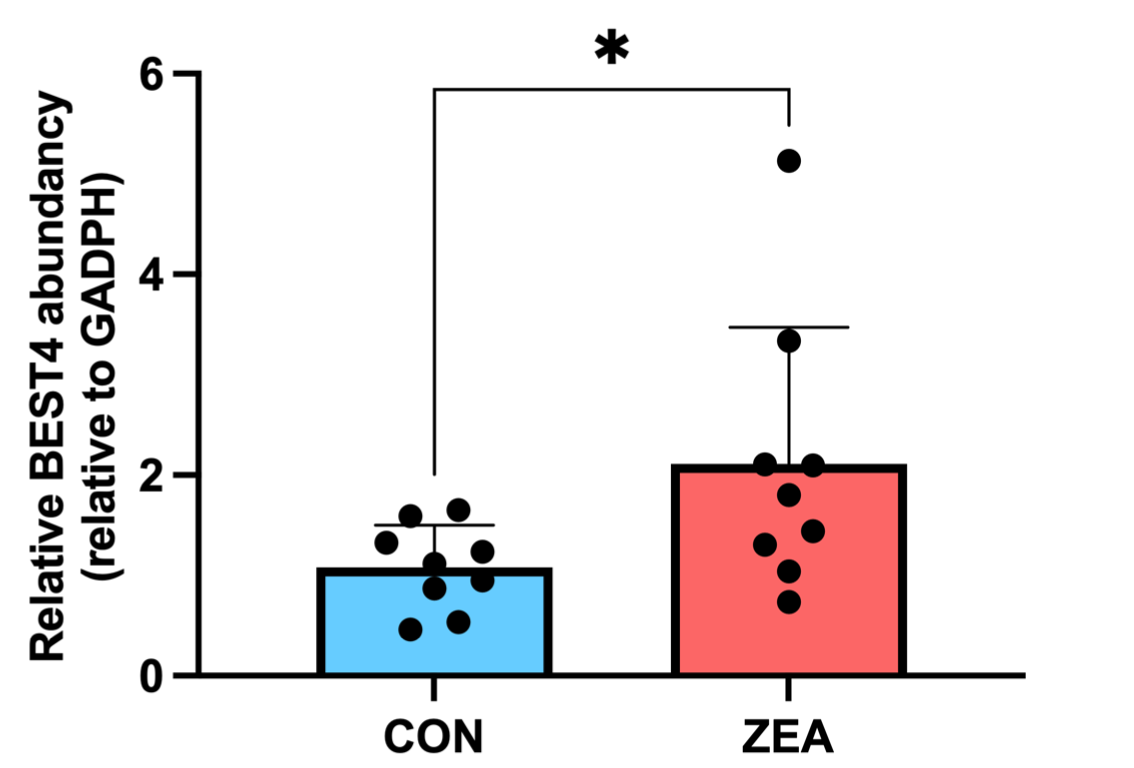

**Fig. S7.** qPCR analysis of the mRNA expression of BEST4. Results shown are mean ± SD, n=9. * p<0.05, * *p<0.01, compared to control group.

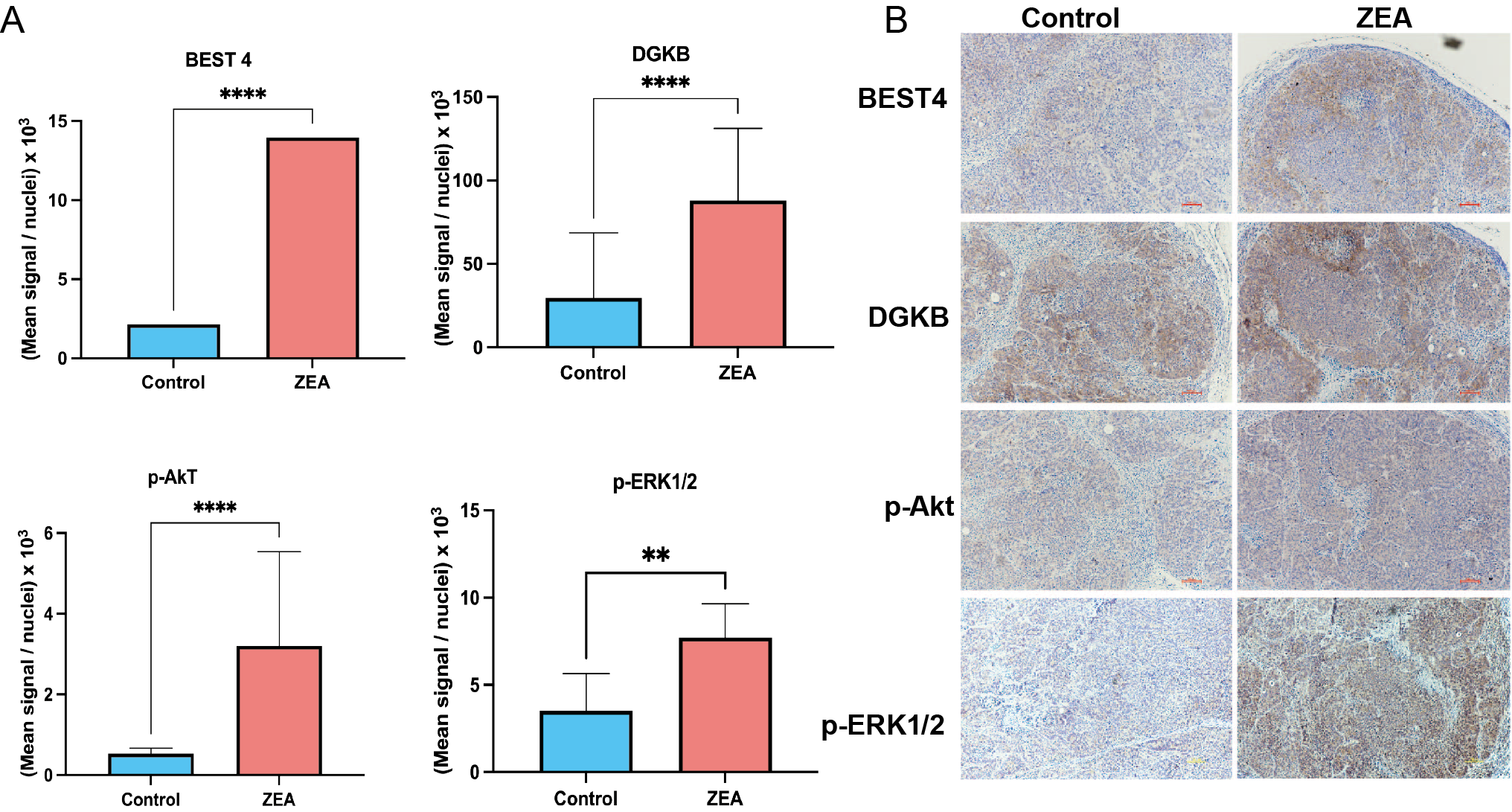

**Fig. S8.** Transcriptional changes in tumor cells in response to ZEA treatment. (A-B) Representative IHC image showing BEST4, DGKB, p-AKT, p-ERK1/2 (100x) and its semi-quantitative analysis. The results shown are mean ± SEM, n=7-8. **p*<0.05, ***p*<0.01, compared to control.

**Table S1.** Amino acids that were altered by ZEA treatment compared to control group.

|  | No. | Amino Acid | ng/g of compound (mean ± SEM) | | *P* value* | Log2 fold change |
| --- | --- | --- | --- | --- | --- | --- |
|  |  |  | Control (n = 6) | ZEA (n = 6) |  |  |
|  | 1 | Phosphoethanolamine | 160.0 ± 53.3 | 79.6 ± 13.6 | 0.01 | -1.00 |
|  | 2 | Taurine | 5242.2.1 ± 157.3 | 2867.4.6 ± 663.2 | 0.02 | -0.86 |
|  | 3 | 1-Methyl Histidine | 32.2 ± 6.5 | 18.3 ± 3.3 | <0.001 | -0.81 |
|  | 4 | Kynurenine | 4.7 ± 0.5 | 2.7 ± 0.3 | <0.001 | -0.81 |
|  | 5 | Aspartic Acid | 58.2 ± 13.1 | 33.6 ± 10.7 | 0.01 | -0.79 |
|  | 6 | 3-Methyl Histidine | 11.0 ± 1.5 | 6.7 ± 1.1 | <0.001 | -0.71 |
|  | 7 | β-Amniobutyric Acid | 0.5 ± 0.2 | 0.3 ± 0.1 | 0.04 | -0.71 |
|  | 8 | Arginine | 634.6 ± 243.3 | 400.9 ± 142.9 | 0.04 | -0.67 |
|  | 9 | Cystathionine | 16.0 ± 4.8 | 10.3 ± 1.7 | 0.04 | -0.64 |
|  | 10 | β-Alanine | 50.5 ± 2.5 | 33.2 ± 4.6 | <0.001 | -0.60 |
|  | 11 | Histidine | 644.0 ± 206.2 | 433.6 ± 85.3 | 0.04 | -0.58 |
|  | 12 | Anserine | 49.2 ± 7.8 | 34.4 ± 3.1 | <0.001 | -0.51 |
|  | 13 | Hydroxylysine | 7.5 ± 1.9 | 5.3 ± 0.7 | 0.02 | -0.49 |
|  | 14 | Ethanolamine | 87.4 ± 15.0 | 64.8 ± 10.9 | <0.001 | -0.43 |
|  | 15 | Phenylalanine | 523.2 ± 92.1 | 386.7 ± 48.9 | 0.01 | -0.43 |
|  | 16 | Citrulline | 489.5 ± 87.9 | 377.4 ± 75.2 | 0.04 | -0.38 |
|  | 17 | Glycine | 1158.2 ± 221.4 | 894.2 ± 150.7 | 0.01 | -0.38 |
|  | 18 | Glutamine | 4090.7 ± 974.1 | 3183.0 ± 416.0 | 0.05 | -0.36 |
|  | 19 | Hydroxproline | 110.8 ± 15.8 | 87.6 ± 24.1 | 0.02 | -0.34 |

*With statistical difference for p < 0.05, Generated by GraphPad Prism 9.0.0. using two-tailed Student’s t-test.

**Table S2.** List of metabolites detected in serum samples of control and experimental groups using Progenesis QI. All the metabolites are presented with fold change (ZEA/Control) and P value.

| **No.** | **Compounds** | **m/z** | **Polarity (+/-)** | **Log2 fold change** | ***P* value*** |
| --- | --- | --- | --- | --- | --- |
| 1 | Coumarinic acid | 165.0547 | + | 1.80 | <0.001 |
| 2 | Uric acid | 169.0359 | + | 1.77 | <0.001 |
| 3 | Malic acid | 133.0141 | - | 1.28 | <0.001 |
| 4 | 2-Diethylaminoethanol | 118.1226 | + | 1.15 | <0.001 |
| 5 | Sinapic acid | 223.0615 | - | 0.97 | <0.001 |
| 6 | Citric acid | 191.0198 | - | 0.94 | <0.001 |
| 7 | Indoleacetaldehyde | 160.0757 | + | 0.89 | <0.001 |
| 8 | Serotonin | 177.1026 | + | 0.87 | <0.001 |
| 9 | Indoleacrylic acid | 188.0708 | + | 0.74 | <0.001 |
| 10 | p-Cresol sulfate | 187.0073 | - | 0.73 | 0.001 |
| 11 | Indole-3-carboxaldehyde | 146.0602 | + | 0.71 | 0.001 |
| 12 | Hippuric acid | 180.0657 | + | 0.64 | 0.001 |
| 13 | Succinylacetone | 203.0829 | - | 0.61 | <0.001 |
| 14 | Indoline | 120.0806 | + | 0.30 | 0.046 |
| 15 | Sphinganine-1-phosphate | 380.2570 | - | -0.35 | 0.008 |
| 16 | Dictyoquinazol A | 357.1096 | - | -0.43 | <0.001 |
| 17 | PC (16:1/20:1) | 830.5914 | - | -0.51 | <0.001 |
| 18 | Hydroxydocosahexaenoic acid (HDoHE) | 343.2270 | - | -0.54 | 0.044 |
| 19 | LysoPE (16:0/0:0) | 436.2836 | - | -0.76 | 0.004 |
| 20 | Hydroxyeicosatetraenoic acids (HETE) | 319.2283 | - | -1.66 | 0.002 |
| 21 | 2-Hydroxycinnamic acid | 165.0547 | + | -2.05 | <0.001 |

*With statistical difference for p < 0.05, Generated by GraphPad Prism 9.0.0. using two-tailed Student’s t-test.

**Table S3.** Metabolites strongly related to tumor weight with spearman's rank correlation coefficient (cutoff: |correlation coefficient|>0.6 and p-value <0.05)

| Metabolite | Correlation coefficient | p value |
| --- | --- | --- |
| Serotonin | 0.747 | 0.005 |
| Uric acid | 0.719 | 0.008 |
| p-Cresol sulfate | 0.705 | 0.01 |
| Indoleacetaldehyde | 0.646 | 0.023 |
| Hippuric acid | 0.607 | 0.036 |
| Succinylacetone | 0.604 | 0.038 |
| Dictyoquinazol A | -0.6 | 0.039 |
| 2-Hydroxycinnamic acid | -0.632 | 0.028 |
| 1-Methyl Histidine | -0.642 | 0.024 |
| Hydroxydocosahexaenoic acid (HDoHE) | -0.642 | 0.024 |
| Sphinganine-1-phosphate | -0.646 | 0.023 |
| β-Alanine | -0.649 | 0.022 |
| Hydroxyeicosatetraenoic acids (HETE) | -0.72 | 0.011 |

**Table S4.** Pearson correlation coefficient and significance level for DA metabolites, DE genes and DA genera (cutoff: |correlation coefficient|>0.8 and FDR <0.05)

|  | Correlation coefficient | p value | FDR |
| --- | --- | --- | --- |
| DGKB and Uric acid | 0.909 | <0.001 | 0.013 |
| ENTPD2 and Coumarinic acid | 0.909 | <0.001 | 0.013 |
| DGKB and p-Cresol sulfate | 0.902 | <0.001 | 0.018 |
| DGKB and Sinapic acid | 0.902 | <0.001 | 0.018 |
| EBF2 and Indoleacetaldehyde | 0.902 | <0.001 | 0.018 |
| BEST4 and Hippuric acid | 0.895 | <0.001 | 0.025 |
| CYP4F29P and Hippuric acid | 0.888 | <0.001 | 0.034 |
| JAK3 and Hippuric acid | 0.888 | <0.001 | 0.034 |
| JAK3 and Coumarinic acid | 0.881 | <0.001 | 0.046 |
| Rikenella and 2-Hydroxycinnamic acid | 0.881 | <0.001 | 0.046 |
| ROBO3 and Hippuric acid | 0.881 | <0.001 | 0.046 |
| Rikenella and 1 Methyl Histidine | 0.874 | <0.001 | 0.029 |
| BEST4 and Tuzzerella | -0.860 | <0.001 | 0.047 |
| PTGIR and Phosphoethanolamine | -0.860 | <0.001 | 0.047 |
| ROBO3 and Beta Alanine | -0.860 | <0.001 | 0.047 |
| ROBO3 and Phosphoethanolamine | -0.860 | <0.001 | 0.047 |
| UCG.005 and Taurine | -0.860 | <0.001 | 0.047 |
| NFATC4 and Glycine | -0.867 | <0.001 | 0.037 |
| UCG.005 and Phosphoethanolamine | -0.867 | <0.001 | 0.037 |
| LINC02593 and Beta Alanine | -0.874 | <0.001 | 0.029 |
| C4orf46 and 2-Diethylaminoethanol | -0.881 | <0.001 | 0.046 |
| C4orf46 and Indole-3-carboxaldehyde | -0.881 | <0.001 | 0.046 |
| DGKB and Tuzzerella | -0.881 | <0.001 | 0.046 |
| ENTPD2 and Ethanolamine | -0.881 | <0.001 | 0.022 |
| ROBO3 and Hydroxyeicosatetraenoic acids (HETE) | -0.881 | <0.001 | 0.046 |
| TMEM167A and Indole-3-carboxaldehyde | -0.881 | <0.001 | 0.046 |
| CYP4F29P and Hydroxyeicosatetraenoic acids (HETE) | -0.888 | <0.001 | 0.034 |
| Paludicola and Succinylacetone | -0.888 | <0.001 | 0.034 |
| PTGIR and Rikenella | -0.902 | <0.001 | 0.009 |
| ENTPD2 and Anserine | -0.916 | <0.001 | 0.004 |
| ENTPD2 and Glycine | -0.930 | <0.001 | 0.002 |
| Rikenellaceae.RC9.gut.group and Serotonin | -0.937 | <0.001 | 0.002 |
| Tuzzerella and Uric acid | -0.937 | <0.001 | 0.002 |

### Supplementary methods

**RNA extraction and Real time reverse transcription polymerase chain reaction (RT-qPCR)**

Total RNA was extracted with illustrate RNAspin Mini RNA Isolation Kit (GE Healthcare Life Sciences; Buckinghamshire, UK) according to the manufacturer’s instructions. The purity of RNA concentration was determined with a NanoDrop 2000 spectrophotometer (Thermo Scientific, DE, USA). cDNA was synthesized using HiScipt II Q-RT SuperMix for qPCR (+ qDNA wiper) (Vazyme, Nanjing, China) according to the manufacturer’s instructions. Quantitative real time PCR (qPCR) was conducted using AceQ qPCR SYBR Green Master Mix (Vazyme, Nanjing, China). The sequence of the primers were as follow: BEST4, 5’-AGAGAAGGTCGGTTGCTTGG-3’ (sense) and 5’ CACCTGGGCTCTAGTCCTCT-3’ (antisense); GADPH, 5’-GACAGTCAGCCGCATCTTCT-3’ (sense) and 5’-GCGCCCAATACGACCAAATC-3’ (antisense). All cDNA samples were run on StepOnePlus Real Time PCR system (Applied Biosystems, CA, USA). The relative RNA expressions were normalized against GADPH using 2^-∆∆CT^ method.
